# Cell cycle-dependent binding between Cyclin B1 and Cdk1 revealed by time-resolved Fluorescence Correlation Spectroscopy

**DOI:** 10.1101/2022.02.22.481435

**Authors:** Martina Barbiero, Luca Cirillo, Sapthaswaran Veerapathiran, Catherine Coates, Camilla Ruffilli, Jonathon Pines

## Abstract

Measuring the dynamics with which the regulatory complexes of the cell cycle machinery assemble and disassemble is a crucial barrier to our understanding that until now has been difficult to address. This considerable gap in our understanding is due to the difficulty of reconciling biochemical assays with single cell-based techniques, but recent advances in microscopy and gene editing techniques now enable the measurement of protein-protein interaction kinetics in living cells.

Here, we apply Fluorescence Correlation Spectroscopy (FCS) and Fluorescence Cross-Correlation Spectroscopy (FCCS) to study the dynamics of the cell cycle machinery, beginning with Cyclin B1 and its binding to its partner kinase Cdk1 that together form the major mitotic kinase. Although Cyclin B1 and Cdk1 are known to bind with high affinity, our results reveal that in living cells there is a pool of Cyclin B1 that is not bound to Cdk1. Furthermore, we provide evidence that the affinity of Cyclin B1 for Cdk1 increases during the cell cycle, indicating that the assembly of the complex is a regulated step. Our work lays the groundwork for studying the kinetics of protein complex assembly and disassembly during the cell cycle in living cells.

## Introduction

The cell cycle relies on the rapid formation and disassembly of regulatory protein complexes. At the core of the cell cycle machinery are the family of Cyclin Dependent Kinases (Cdks – reviewed in Morgan, 1997) that are activated by binding a cyclin subunit. Cyclin B1 binds and activates Cdk1 to form the major mitotic kinase that is required for cells to enter mitosis (Strauss et al., 2017). The Cyclin B1-Cdk1 complex itself binds to an accessory Cks protein (Cyclin-dependent kinases regulatory subunit Cks1 or Cks2) that recognises and binds to phospho-threonine (Brown et al., 2015; Draetta et al., 1987; Kõivomägi et al., 2013, 2011; Patra et al., 1999; Richardson et al., 1990; reviewed in Pines, 1996). Cyclin B1 levels start to increase late in S phase and it continues to accumulate in the cytoplasm of G2 cells (Pines and Hunter, 1991, 1989). The activation of the Cyclin B1-Cdk1-Cks complex sets the time for mitotic entry in all eukaryotes studied to date (reviewed in Nurse, 1990), whereas its inactivation at metaphase via the ubiquitin-mediated proteolysis of Cyclin B1 is required for cells to exit mitosis (Clute and Pines, 1999; Murray et al., 1989; Murray and Kirschner, 1989).

At mitosis, the entire cell architecture is reorganised in a matter of minutes. Biochemical analyses have shown that the mitotic kinases and phosphatases have multiple substrates and are often components of several different complexes. Live cell analyses have shown that cell cycle regulators are often highly dynamic; for example, Fluorescence Recovery After Photobleaching shows that Cyclin B1 fluxes rapidly on and off the mitotic spindle (Richardson and Pines, unpublished results). Similarly, Förster Resonance Energy Transfer probes specific for different components reveal spatial gradients of activity (reviewed in Pines and Hagan, 2011). Thus, to understand how the cell cycle machinery works, we must measure the kinetics with which regulatory complexes assemble and disassemble, both with respect to cell cycle time and position in the cell.

Biophysical methods such as X-ray crystallography and electron microscopy provide important structural information, and biochemical assays have proved to be invaluable for the understanding the underlying biochemical properties, but because they require populations of cells, they have very limited temporal and spatial resolution, and cannot measure protein dynamics and interactions i*n vivo*. By contrast, Fluorescence correlation spectroscopy (FCS) accurately estimates the *in vivo* concentration and dynamics of fluorescently tagged molecules with high spatial and temporal resolution by analysing the intensity fluctuations in a confocal volume (femtoliter scale) (Brock and Jovin, 1998; Enderlein et al., 2005; Kim et al., 2007, p. 200; Krichevsky and Bonnet, 2002; Magde et al., 1974; Yu et al., 2021). Furthermore, fluorescence cross-correlation spectroscopy (FCCS) can measure the interaction between two biomolecules tagged with spectrally different fluorophores (Ries et al., 2009; Schwille et al., 1997; reviewed in Bacia et al., 2006). Until recently, two factors have limited the application of FCS and FCCS in living cells: the difficulty of expressing a fluorescently labelled protein at a physiologically relevant concentration (i.e. not by overexpression), and the presence of the unlabelled version of the same protein in the cell. The advent of CRISPR/Cas9 gene editing now enables a fluorescent tag to be incorporated into both alleles of a gene to produce a uniformly labelled protein population (reviewed in Doudna and Charpentier, 2014). Thus, a combination of FCS imaging with CRISPR/Cas9 gene editing offers a platform to study the rapid dynamics of protein complex assembly and disassembly that drive the cell cycle *in vivo*.

In this work, we use CRISPR/Cas9 to tag Cyclin B1 biallelically with the fluorescent protein mEmerald (Cubitt et al., 1998) in untransformed human Retinal Pigment Epithelium (RPE – hTERT, hereafter referred to as RPE) cells, to perform FCS and FCCS measurements through the cell cycle to analyse the dynamics of its assembly into active complexes. Our FCS analysis reveals the existence of two species of Cyclin B1 of different molecular sizes, consistent with a population of free CyclinB1 and a population of CyclinB1 bound to its interacting kinase Cdk1. We have validated these results by immunodepletion of Cdk1 in RPE lysates, which confirms the existence of a pool of Cyclin B1 not bound to Cdk1. FCS and FCCS measurements reveal that the fraction of Cyclin B1 bound to Cdk1 increases as cells progress through G2 phase and this is explained by an increase in the affinity of binding. We conclude that the binding between Cyclin B1 and Cdk1 is cell cycle regulated. Overall, our results demonstrate that FCS and FCCS can be used to measure the concentration and interactions of cell cycle proteins in living cells in a time-resolved manner, which will increase our understanding of how the cell cycle is regulated with such precision.

## Results

### There are two populations of Cyclin B1 in cells

FCS is an imaging-based technology that relies on fluorescence measurements to estimate a diffusion coefficient. To apply FCS to Cyclin B1 we used CRISPR/Cas9^D10A^ nickase (Shen et al., 2014) to introduce the mEmerald coding sequence at the 3’ of the CCNB1 open reading frame. Biallelic-tagged clones were identified by PCR and immunoblot analysis (Fig. S1A). In agreement with previous reports (Clute and Pines, 1999; Hagting et al., 1998; Jackman et al., 2020, 2003), Cyclin B1-mEmerald localised to the cytoplasm, particularly the centrosomes, of interphase cells and was recruited to the spindle, the chromosomes, and the spindle poles of mitotic cells (Fig S1B, C). The mitotic timing, Spindle Assembly Checkpoint (SAC) response and chromosome number of the RPE CCNB1-mEmerald^+/+^ cells did not significantly differ from the parental cell line (Fig S1D, 1E), indicating that the addition of mEmerald did not affect Cyclin B1 function and that we could use the fusion protein to report on the proper behaviour of Cyclin B1.

The autocorrelation functions (ACF) from FCS measurements on Cyclin B1-mEmerald fitted better to a 3D two-particle triplet model (3D-2p-triplet) than a 3D one-particle triplet model (3D-1p-triplet) (See Materials and Methods), in both in the cytoplasm and the nucleus (Fig. 1A-C). This indicated that two populations of Cyclin B1-mEmerald existed: a fast-diffusing fraction with a diffusion coefficient (D) of ~ 35 μm^2^/s; and a slow-diffusing fraction with a D of ~ 8 μm^2^/s (Figure 1D).

**Figure 1:**
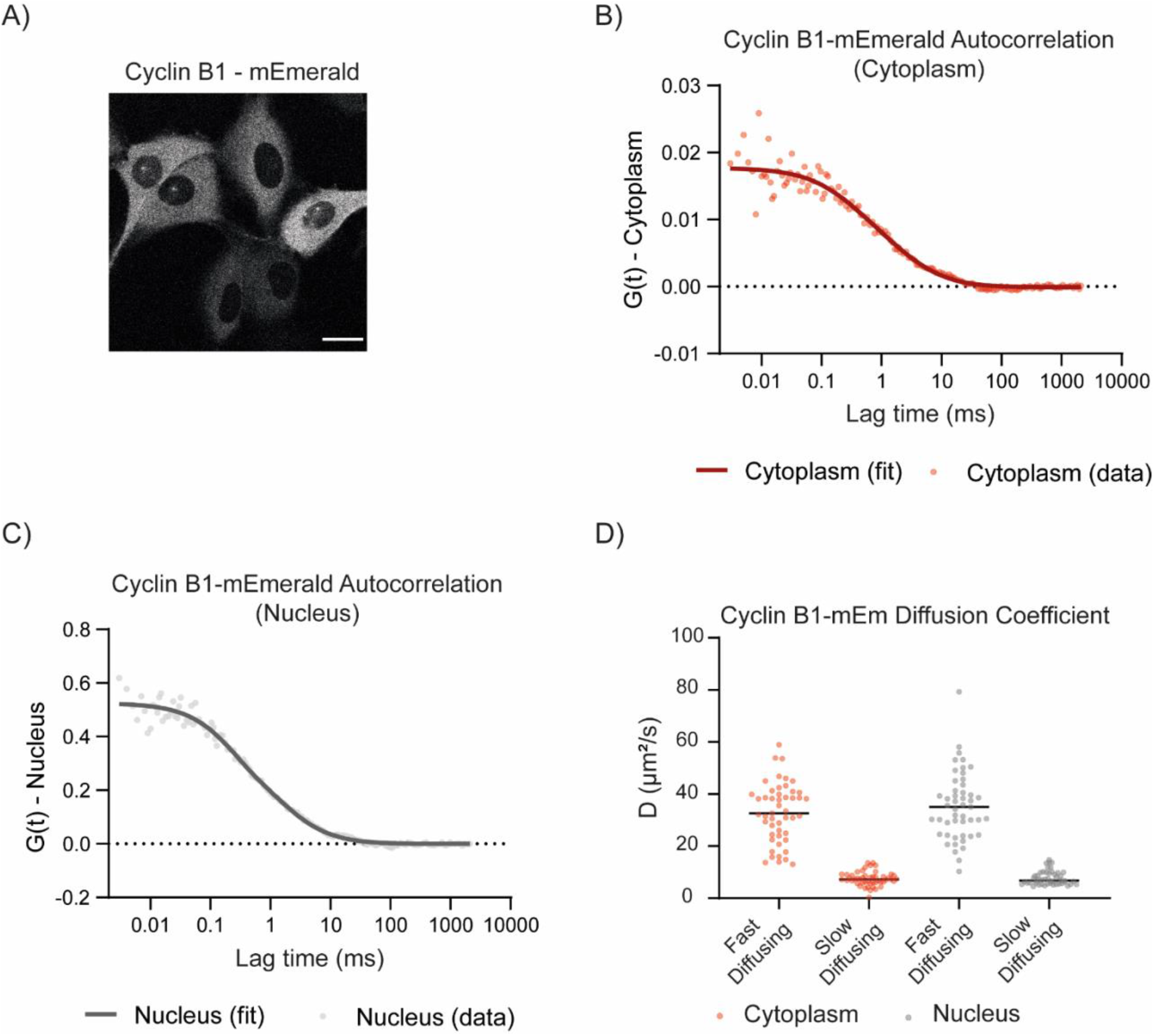
Cyclin B1 size in RPE cells. A) Representative fluorescence confocal image of CCNB1-mEmerald^+/+^ cells. Scale bar corresponds to 20 μm. B) Graph representing the autocorrelation function of Cyclin B1-mEmerald over time in the cytoplasm C) Graph representing the autocorrelation function of Cyclin B1-mEmerald over time in the nucleus D) Dot plot representing the diffusion coefficient of Cyclin B1-mEmerald species. Horizontal black lines represent median values. For all experiments N= 3 or more.

The apparent sizes of the two Cyclin B1-mEmerald species can be estimated using the Stokes-Einstein equation (Equation 1) provided the viscosity of RPE cells at 37 °C is known.

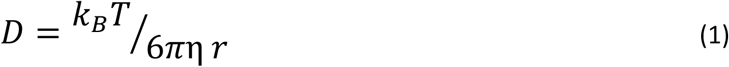

To calculate the viscosity of RPE cells, we used a RPE cell line stably expressing GFP and measured its D in the cytoplasm and nucleus using FCS (Fig. S2A, B). Fitting the ACFs with a 3D one-particle triplet (3D-1p-triplet) model, we obtained a D of 42 ± 5 μm^2^/s for GFP in RPE cells at 37°C. The hydrodynamic radius of GFP was previously reported to be ~ 2 nm (Hink et al., 2000; Luo et al., 2019); therefore, we estimated that the mean viscosity of RPE cells at 37°C was 2.4 ± 0.7 mPa.s in the cytoplasm and 2.6 ± 0.6 mPa.s in the nucleus (Fig. S2C). We obtained comparable results using the mVenus fluorescent protein (Fig. S2D). Using these values in equation 1 gave the hydrodynamic radius of the fast-diffusing Cyclin B1-mEmerald species as 3-4 nm and the slow diffusing Cyclin B1-mEmerald species as 8-11 nm. The hydrodynamic radius of Cyclin B1-mEmerald estimated from the structures of the proteins is ~ 4 nm, whereas that of the Cyclin B1-mEmerald-Cdk1-Cks complex is ~ 7.5 nm (Brown et al., 2015; Fleming and Fleming, 2018). These values are in the range of the hydrodynamic radii of the two populations that we measured by FCS. Therefore, we conclude that the two populations of Cyclin B1 are an unbound freely diffusing monomer and a fraction of Cyclin B1 bound to Cdk1 and Cks.

To test our conclusion, we used quantitative immunoblotting to assay for Cyclin B1 in cell lysates after immunodepleting Cdk1. Immunodepleting Cdk1 should remove all its bound Cyclin B1 because Cyclin B1 binds Cdk1 with high affinity (Brown et al., 2015; Desai et al., 1995). Two sequential immunodepletions of Cdk1 in G2 phase RPE CCNB1-mEmerald^+/+^ cells reduced Cdk1 to 44.0 (± 11.3)% and 6.0 (± 1.4)% (Mean ± SD) of its original levels (Fig 2A – compare lane 1 with lane 2 and 3), whereas Cyclin B1 levels dropped to 58.0 (± 18.4)% and 29.0 (± 2.8)% of its original levels (Fig 2A – compare lane 1 with lane 2 and 3). This indicated that approximately 18% of Cyclin B1 was not bound to Cdk1 (Fig 2C). We obtained similar results using parental RPE cells, excluding the possibility that tagging Cyclin B1 affected its binding to Cdk1 (Fig. 2C, S3A, B). We observed a Cyclin B1 signal even after depleting Cdk1 below the detection threshold (Fig. 2D, S3C), further demonstrating that some Cyclin B1 did not bind Cdk1. (Note that, as expected, we were unable to detect Cdk2, in immunoprecipitates of Cyclin B1 (Fig. 2E, S3D – compare lane 2 with lane 1 and 3), indicating that Cdk2 did not bind significantly to Cyclin B1 in vivo.) Overall, these data confirmed our conclusions from our FCS data that there were two populations of Cyclin B1 in RPE cells: monomeric Cyclin B1 and the Cyclin B1-Cdk1-Cks complex.

**Figure 2:**
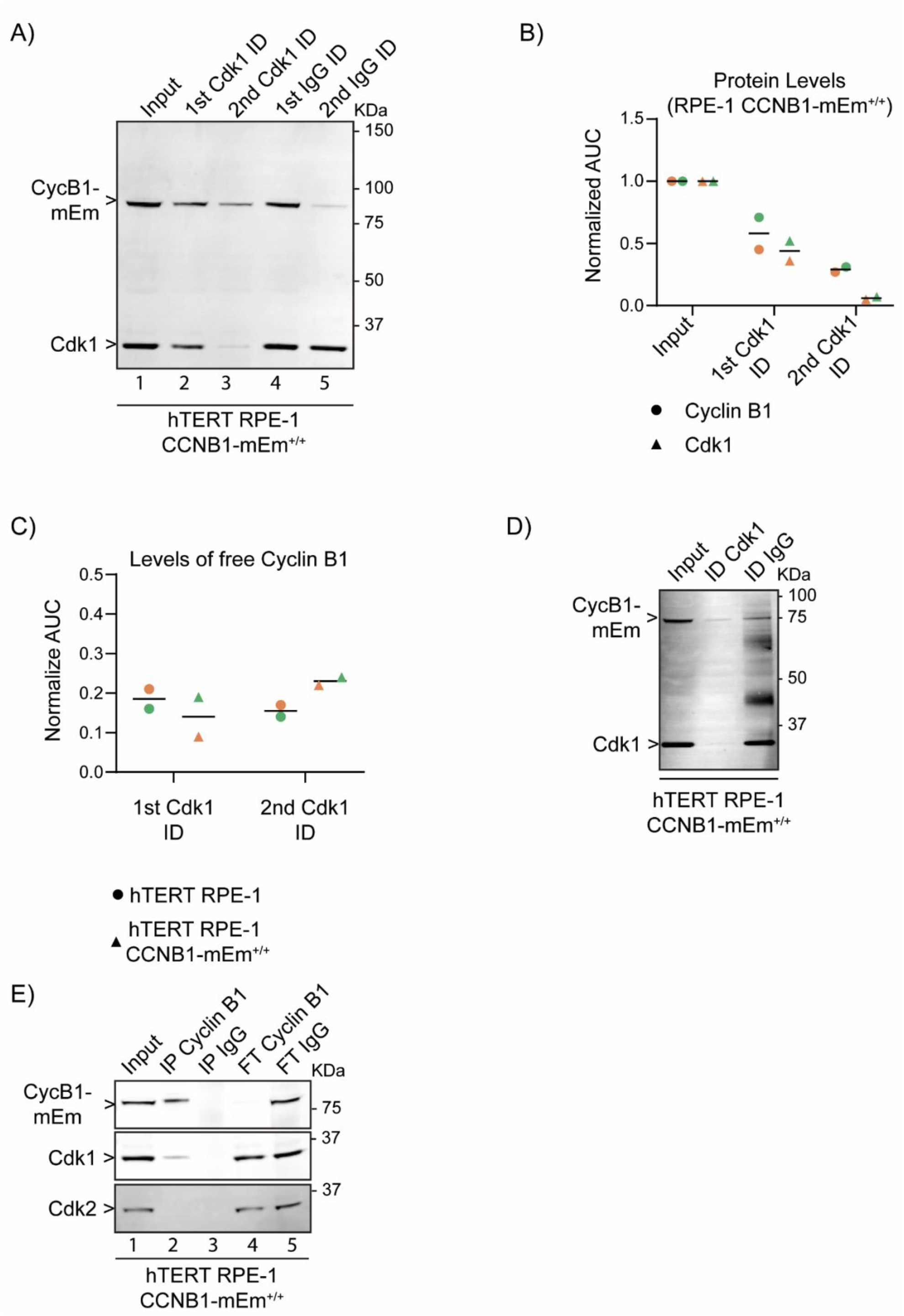
A fraction of Cyclin B1 is not bound to Cdk1 in RPE CCNB1-mEM^+/+^ cells. A) Anti-Cyclin B1 and anti-Cdk1 immunoblot of G2 phase cell lysates before (1^st^) and after (2^nd^ and 3^rd^ lanes) immunodepleting Cdk1, compared with control immunodepletion with IgG (4^th^ and 5^th^ lanes). (Note that we do not have a definitive explanation for the loss of of Cyclin B1 on control beads in the 2^nd^ depletion. This was not consistent, compare Supplemental Figure 3, panels A and C.) B) Quantification of Cyclin B1 and Cdk1 levels before and after immunodepletion of Cdk1. C) Quantification of Cyclin B1 levels before and after immunodepletion of Cdk1 from parental RPE1 cells and from RPE-CCNB1-mEmerald cells. D) Anti-Cyclin B1 and anti-Cdk1 immunoblot of G2 phase cell lysates before and after immunodepletion of Cdk1 to at or below detection levels, or after control immunodepletion. E) Anti-Cyclin B1, anti-Cdk1 and anti-Cdk2 immunoblots of G2 phase cell lysates before (1^st^ lane) and after immunoprecipitation with anti-Cyclin B1 antibody (2^nd^ lane), or immunoprecipitation with anti-IgG control antibody (3^rd^ lane) and the unbound fractions from the respective immunodepletions (4^th^ and 5^th^ lanes). For all graphs, individual dots represent biological replicates, horizontal lines indicate median values. AUC = area under the curve. Molecular mass indicated for all gels on the right. For all panels, N=2 independent experiments. ID= immunodepletion, IP = immunoprecipitation, FT = Flow Through, CycB1-mEm = Cyclin B1-mEmerald.

### Cyclin B1-Cdk1 interaction can be measured using FCCS

Our biochemical data indicated that FCS measurements had accurately identified two populations of Cyclin B1 in living cells; therefore we should be able to detect the Cyclin B1-Cdk1 interaction *in vivo* using FCCS. With this aim we generated a RPE cell line expressing a mEmerald-mScarlet fusion protein as a positive control for FCCS, and introduced mScarlet alone into the RPE CCNB1-mEmerald^+/+^ cell line as a negative control (Bindels et al., 2017). In the cells expressing the mEmerald-mScarlet fusion protein we obtained an ACF for each fluorophore, plus a cross-correlation function with a cross-correlation quotient *q* of ~ 55%-65% (Fig. S4A). That the cross-correlation quotient *q* was less than 100% is explained by incomplete maturation and extended dark state residence of red fluorescent proteins (Balleza et al., 2018; Dunsing et al., 2018; Foo et al., 2012; Hillesheim et al., 2006). In the negative control RPE CCNB1-mEmerald^+/+^ cells expressing mScarlet there was no detectable cross-correlation (*q* <5%) between Cyclin B1-mEmerald and mScarlet (Fig. S4B).

Having validated our FCCS measurement, we were in a position to measure the cross-correlation between Cyclin B1 and Cdk1. To enable this, we introduced a tetracycline-inducible construct encoding Cdk1 tagged at the carboxyl terminus with mScarlet (Fig 3A) into the CCNB1-mEmerald^+/+^ cell line. Co-immunoprecipitation followed by immunoblotting showed that Cyclin B1-mEmerald bound to Cdk1-mScarlet (Fig. S4C), and FCCS revealed cross correlation between Cyclin B1-mEmerald and Cdk1-mScarlet (*q* ~25%-35%) (Fig. 3B). We concluded that FCCS could measure protein-protein interactions in living cells.

**Figure 3:**
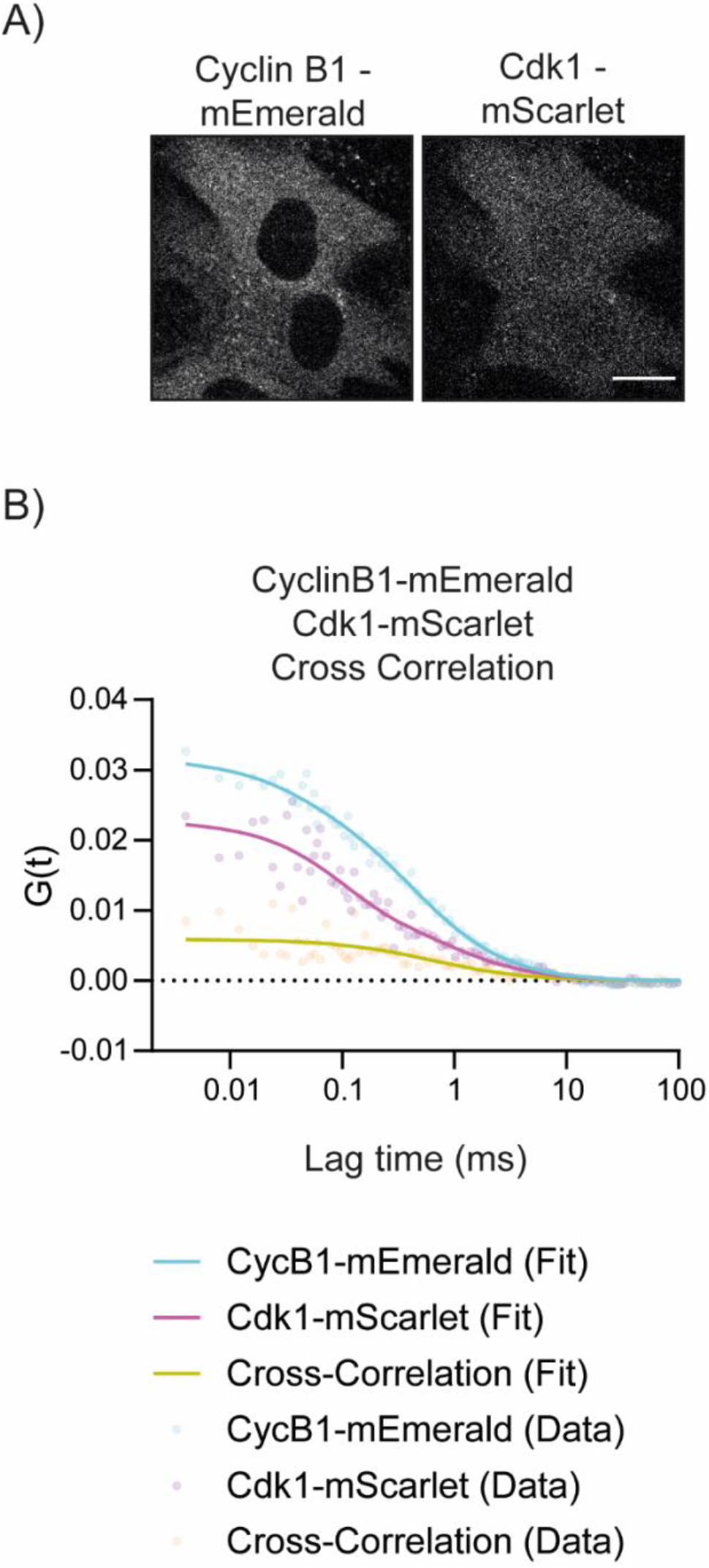
FCCS detects the interaction between Cyclin B1 and Cdk1. A) Representative fluorescence confocal image of RPE CCNB1-mEmerald^+/+^ cells expressing Cdk1-mScarlet from a tetracycline-inducible promoter. Scale bar corresponds to 20 μm. B) Graph of the autocorrelation of Cyclin B1-mEmerald (Cyan), Cdk1-mScarlet (Magenta) and the Cross-Correlation between the two (Yellow). For all panels, N=2 independent experiments.

### Time resolved measurement of Cyclin B1 concentration and complex fraction

Our results revealed that Cyclin B1 existed both as a free monomer and in a complex with Cdk1, and we wondered whether the relative abundance of these two populations changed through the cell cycle. To determine this we synchronized cells in G1 phase using the Cdk4/6 inhibitor Palbociclib (Scott et al., 2020; Trotter and Hagan, 2020) and assayed cells at specific times after release from the arrest. At each time point we fixed and stained the cells for flow cytometry-based cell cycle analysis (Fig. 4A), analysed Cyclin B1 size and concentration through FCS (Fig. 4B, C and Fig S5 A, B), and immunoprecipitated Cyclin B1 from cell lysates to assess its binding to Cdk1 (Fig. 4D). FCS measurements revealed that the concentration Cyclin B1 increased over time following an exponential function (Fig 4C and Fig S5B). We observed a similar Cyclin B1 increase by immuno-blot analysis (Fig. 4D – left panel). FCS analysis also revealed that the fast-diffusing monomeric Cyclin B1 was the dominant fraction until 9 hours after release, after which the slow diffusing Cyclin B1-Cdk1-Cks complex became prevalent (Fig. 4C and Fig S5B). In agreement with this, we observed an increase in the ratio of Cdk1 binding to Cyclin B1 in Cyclin B1 immunoprecipitates at later time points after release (Fig. 4D – right panel, S5C).

**Figure 4:**
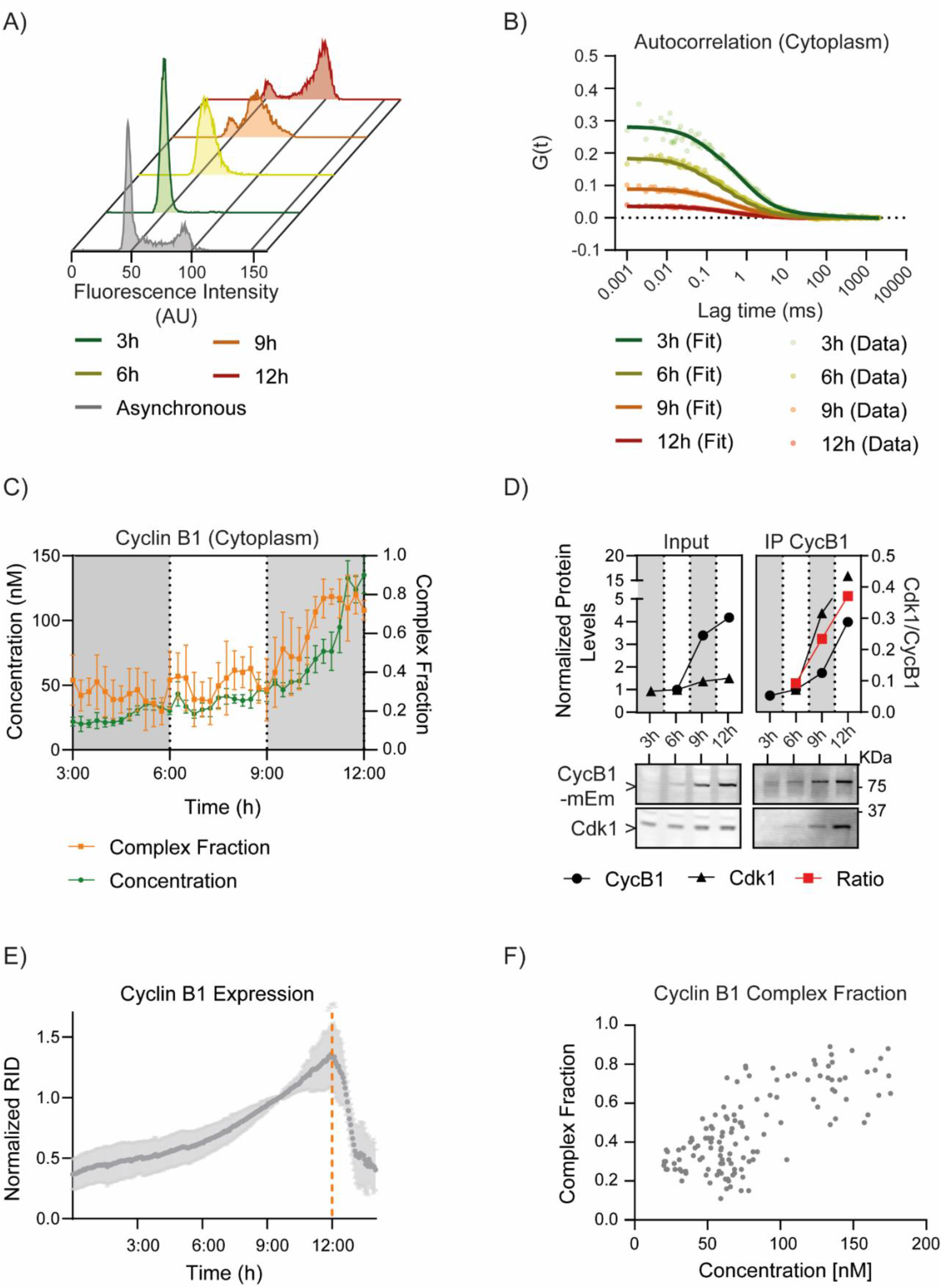
Time resolved measurement of Cyclin B1 size and concentration. A) Flow cytometry profiles of Propidium iodide-stained RPE CCNB1-mEmerald^+/+^ cells: asynchronous and at 3, 6, 9 and 12 hours after release from Palbociclib-arrest. B) Plots of the FCS autocorrelation functions of Cyclin B1-mEmerald at the indicated time points after release from Palbociclib-arrest. Each dot represents the measurement from one cell. C) Graph showing time-resolved Cyclin B1-mEmerald concentration (left axis) and the fraction of Cyclin B1 in complex with Cdk1 (right axis) estimated from FCS measurements in the cytoplasm. D) Top panel: quantification of the protein levels measured from anti-Cyclin B1 and anti-Cdk1 immuno-blots (bottom panel, molecular mass indicated on the right) of either synchronized lysates (left) or Cyclin B1 immunoprecipitates (right). The ratio between Cdk1 and Cyclin B1 is plotted in red on the right axis. E) Quantification of Cyclin B1 fluorescence levels (normalised Raw Integrated Density, RID) over time measured by widefield fluorescence microscopy in unsynchronized RPE CCNB1-mEmerald^+/+^ cells. Orange dotted line indicates nuclear envelope breakdown. F) Quantification of FCS-derived measurements of the fraction of Cyclin B1 bound to Cdk1 over total Cyclin B1 levels, plotted against total Cyclin B1 concentration. In all graphs error bars indicate standard deviation. For all panels, N=2 independent experiments, time indicates hours after Palbociclib release.

To exclude the possibility that treatment with Palbociclib might perturb the normal behaviour of Cyclin B1, we analysed asynchronous cells. The fluorescence level of Cyclin B1-mEmerald increased exponentially before nuclear envelope breakdown (Fig. 4E, S5D). This exponential increase measured by widefield epifluorescence agreed with the exponential increase in Cyclin B1-mEmerald concentration in synchronised cells. FCS measurements in asynchronous cells showed a correlation between Cyclin B1 concentration and the fraction of slow diffusing Cyclin B1 complex in unsynchronized cells: in cells with a low concentration of Cyclin B1 (S phase and early G2 phase), most Cyclin B1 diffused fast, whereas in cells with higher concentrations of Cyclin B1 (mid/late G2 phase) the slowly diffusing Cyclin B1 population was dominant (Fig 4F). This agreed with the data in Fig. 4C. Thus, we concluded that synchronisation with Palbociclib did not perturb the behaviour of Cyclin B1, and that the percentage of Cyclin B1 bound to Cdk1 changed from about 30%-40% in S phase to about 70%-80% in late G2 phase.

### Estimating Cyclin B1-Cdk1 binding affinity in living cells

We wanted to know whether the change in the proportion of Cyclin B1 binding to Cdk1 as cells progressed through G2 phase would fit with a simple model where binding to a constant amount of Cdk1 was driven by an increase in concentration of Cyclin B1. We modelled Cyclin B1 expression with a single exponential function, and fitted Cdk1 to a straight-line equation, using 629 nM as the initial Cdk1 concentration (inferred from Beck et al., 2011) and 28 nM as the Cyclin B1-Cdk1 K_D_ (Dissociation Constant, Brown et al., 2015). We calculated the fraction of Cyclin B1 in complex with Cdk1 according to our model using equation (2).

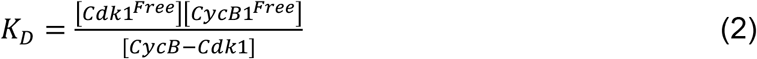

Using these parameters, more than 95% of Cyclin B should have been bound to Cdk1 even at low levels of Cyclin B1, which did not match our experimental data (Fig 4C). The discrepancy could be because the value of the dissociation constant (K_D_) measured *in vitro* might be different *in vivo*, where the conformation and interactions of proteins might vary considerably (reviewed in Lipinski and Hopkins, 2004). This prompted us to use FCCS to measure the K_D_ of the Cyclin B1-Cdk1 complex *in vivo* (See Materials and Methods). We arrested RPE CCNB1-mEmerald^+/+^ cells expressing Cdk1-mScarlet in G1 phase with Palbociclib and measured the K_D_ for Cyclin B1-Cdk1 at different time points following release from the arrest. The synchrony of the cells was assayed in parallel using flow cytometry (Fig. 5A). We measured the K_D_ for cells 6, 9 and 12 hours after release (the low levels of Cyclin B1 prevented us from measuring the K_D_ at 3 hours). We observed the effective K_D_ for Cyclin B1-mEmerald and Cdk1-mScarlet reduced from 270 nM in early G2 phase (6h post release from Palbociclib) to 112 nM just before mitosis (12h post release from Palbociclib) (Figure 5B, C, D). Using 112 nM as the K_D_, our model (equation 2) predicted that 87% of Cyclin B1 should be in complex with Cdk1 at 12 hours post Palbociclib release, in good agreement with the 79% obtained through FCS.

**Figure 5:**
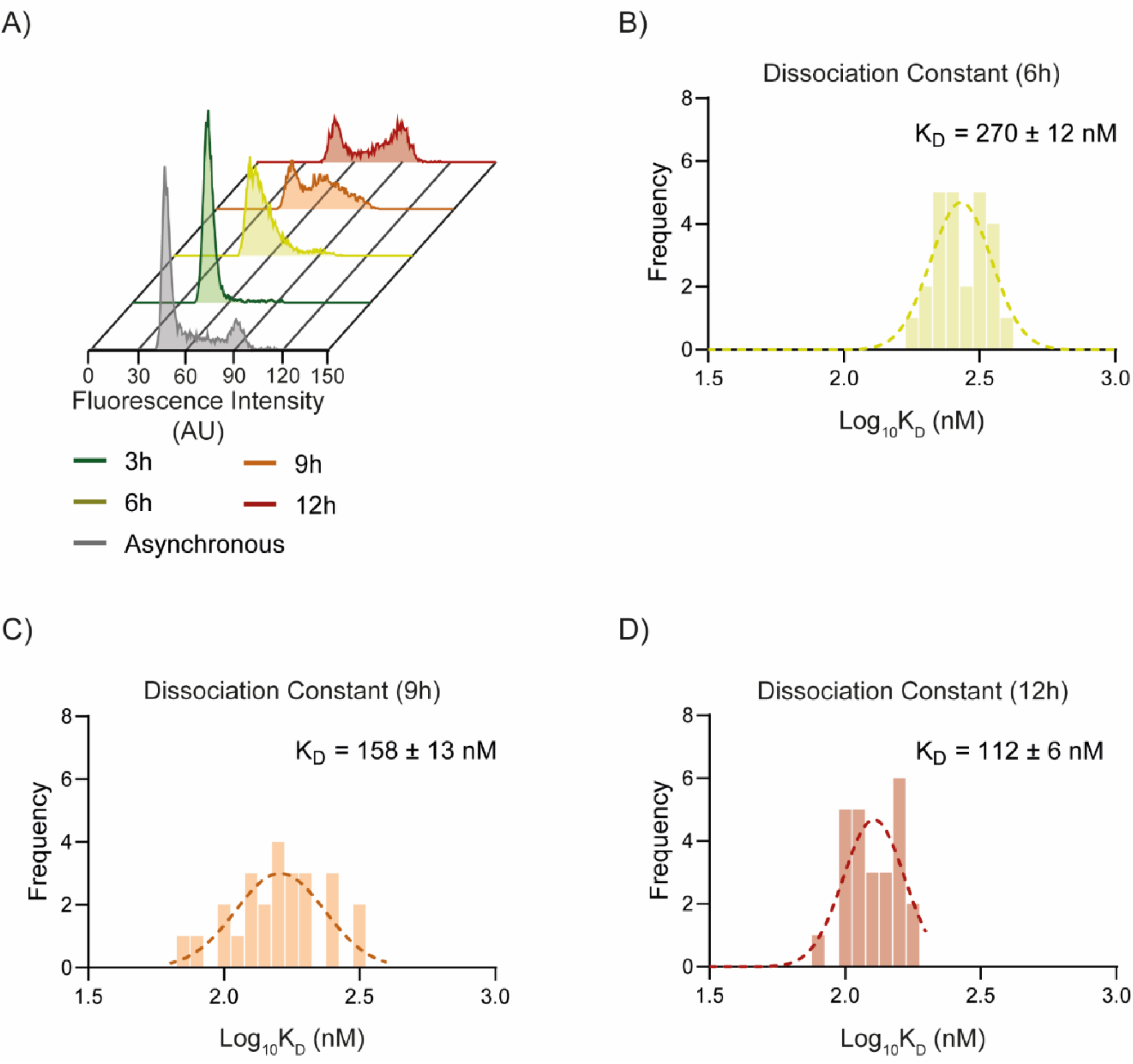
The Dissociation Constant of the Cyclin B1-Cdk1 complex decreases through the cell cycle. A) Flow cytometry profiles of Propidium iodide-stained RPE CCNB1 - mEmerald^+/+^ Cdk1-mScarlet cells: asynchronous and at 3, 6, 9 and 12 hours after release from Palbociclib-arrest. B - D) Frequency histograms of the Log (K_D_) values measured for Cyclin B1-mEmerald and Cdk1-mScarlet by FCCS (see Materials and Methods) at 6 (B), 9 (C), and 12 (D) hours after release from a Palbociclib-arrest. In all histograms, the dotted line represents the Gaussian fit. K_D_ is indicated as mean ± SEM (Standard Error of the Mean). For all panels, N=2 independent experiments.

Altogether, these results demonstrate that the K_D_ value of Cyclin B1-Cdk1 interaction was higher *in vivo* (112 nM) than *in vitro* (28 nM), and that the affinity between Cyclin B1 and Cdk1 increased as cells progressed through G2 phase to peak just before mitosis. This implied that the binding between Cyclin B1 and Cdk1 might be a regulated step.

## Discussion

To understand how the multiple components of the cell cycle machinery coordinate the profound changes in cell architecture with temporal and spatial precision requires that we can measure the assembly and disassembly of regulatory complexes in proliferating cells. FCS and FCCS provide quantitative information on living single cells that combines some of the advantages of biochemistry with those of imaging, such that we can measure protein concentration and protein complex assembly and disassembly in a spatially and temporally defined manner, making them valuable tools to study the rapid events underlying cell cycle progression.

FCS and FCCS have been previously used as tools to study cell cycle; in particular, pioneering work using high-throughput FCS by the Ellenberg lab (Wachsmuth et al., 2015; Walther et al., 2018). Wachsmuth and colleagues measured the temporal changes in diffusion, concentration and cross-correlation ratio of the cell cycle proteins Aurora B and INCENP (Inner Centromere Protein) (Wachsmuth et al., 2015). Walther et al. 2018 used FCS of endogenously-tagged condensins to measure the number of condensin complexes on mitotic chromosomes (Walther et al., 2018). In our study, we additionally measured intracellular viscosity in order to estimate the hydrodynamic radii of Cyclin B1 and Cyclin B1/Cdk1 complex. In addition, we used FCCS to measure *in vivo* the binding affinity between Cyclin B1 and Cdk1.

We estimate that in RPE cells the concentration of Cyclin B1 increases from ~20 nM in S phase to ~150 nM in late G2 phase. Our measurements agree with previous studies (Beck et al., 2011; Karuna et al., 2020; Thomas et al., 2005), although other works reported higher concentrations (Arooz et al., 2000; Frisa and Jacobberger, 2009; Xu and Chang, 2007). The variability is likely explained by differences in the cell lines measured and in the methodology. Indeed, FCS tends to underestimate concentrations due to the complex photophysics, incomplete maturation and fluorescence probability (p_f_) of fluorescent proteins – for example green fluorescent proteins have a pf of ~70-80% and red fluorescent proteins have a pf of ~50-60% (Balleza et al., 2018; Dunsing et al., 2018; Foo et al., 2012; Hillesheim et al., 2006; Schenk et al., 2004). On the other hand, immunoblotting-based measurements are complicated by the requirement for careful calibration of the antibodies and a linear detection method.

Although Cdk1 levels are often considered to be constant during the cell cycle, our quantitative immunoblotting revealed that Cdk1 levels increase by 60% as cell progress through G2 phase (Fig. 4D), in agreement with several other reports (Becher et al., 2018; Ly et al., 2017; Olsen et al., 2010). It is not clear whether newly synthesised Cdk1 has different properties compared to the Cdk1 persisting from the previous cell cycle, but if it does this could offer an explanation for the regulated assembly of Cyclin B1-Cdk1 complexes that our data imply (see below, Welch and Wang, 1992).

Our FCS measurements reveal that Cyclin B1 exists as two distinct species in RPE cells: monomeric Cyclin B1, and Cyclin B1 in complex with its interacting partner Cdk1 (Fig. 1). We validated this result using Cdk1 immunodepletion (Fig. 2) and FCCS with Cdk1-mScarlet (Fig. 3). A pool of monomeric Cyclin B1 is in line with previous conclusions from cell lysates (Arooz et al., 2000; Pines and Hunter, 1989) but to our knowledge our study represents the first measurement of such a pool in intact cells. Using FCS in a time-resolved manner, we observed a sharp increase in the fraction of Cyclin B1 binding to Cdk1 in late G2 phase (Fig. 4), which correlates with an increase in affinity between the two proteins (Fig. 5). The effective K_D_ values we estimate from FCCS are higher than those reported *in vitro* (Brown et al., 2015; Desai et al., 1995) but this can be explained by a) the molecular crowding of the cytoplasm compared to the environment found in a test tube; b) the photophysics of the fluorophores influencing FCCS measurements; and c) the competition for Cyclin B1 between endogenous Cdk1 and Cdk1-mScarlet (Foo et al., 2012). Nevertheless, the increase in affinity between Cyclin B1 and Cdk1 as cells progress through the cell cycle implies some regulation of the binding dynamics of the two proteins. The nature of this regulation is as yet undefined, but it is likely to be one or more post-translational modifications. A potential mechanistic explanation is through modulation of the phosphorylation of Cdk1 on its T-loop (T161), which favours Cdk1-Cyclin B1 binding (Ducommun et al., 1991; Gould et al., 1991; Larochelle et al., 2007; Lee et al., 1999; Norbury et al., 1991). T161 is phosphorylated by CAK (Cdk1 Activating Kinase); however the major CAK in vertebrate cells – a trimeric protein complex constituted by Cdk7, Cyclin H and MAT1 (Ménage à trois 1) - is constitutively active during the cell cycle (Darbon et al., 1994; Fesquet et al., 1993; Fisher and Morgan, 1994; Larochelle et al., 2007; Poon et al., 1993; Solomon et al., 1993; Tassan et al., 1994). There is evidence for an alternative CAK that recognises Cdk2 (Kaldis et al., 2001; Liu et al., 2004), whether this supplemental CAK activity is cell cycle regulated and could also recognise Cdk1 remains to be investigated. Conversely, the phosphatases that remove T-loop phosphorylation could be regulated. The T-loop of monomeric Cdk2 is dephosphorylated by phosphatase 2C and CDKN3 (Cyclin-dependent kinase inhibitor 3; formerly known as KAP - Kinase-associated phosphatase) (Cheng et al., 2000; Gyuris et al., 1993; Hannon et al., 1994; Poon and Hunter, 1995), but their role in regulating Cdk1 and cell cycle regulation is less clear (Nalepa et al., 2013; Smedt et al., 2002). Alternatively, Coulovan and colleagues reported coupling between the phosphorylation of T14 and T161 on Cdk1, whereby regulation of T14 phosphorylation influenced T161 phosphorylation and interaction with Cyclin B1 (Coulonval et al., 2011). Aside from phosphorylation, K-33 acetylation also appears to affect the interaction between Cdk1 and Cyclin B1, and may be subjected to cell cycle regulation (Deota et al., 2019).

In conclusion, we have established the means to measure the kinetics with which protein complexes assemble and disassemble in living cells, and our data reveal a previously unsuspected regulated step in the entry to mitosis where the assembly of the major mitotic kinase, Cyclin B1 and Cdk1, increases as cells progress through G2 phase.

## Material and Methods

### Cell culture and synchronisation

hTERT RPE-1 FRT/TO cells were cultured in F12/DMEM (Sigma-Aldrich) medium supplemented with GlutaMAX (Invitrogen), 10% FBS (Gibco), 0.348% sodium bicarbonate, penicillin (100 U/ml), streptomycin (100 μg/ml), and Fungizone (0.5 μg/ml). Cells were maintained in a humidified incubator at 37°C and 5% CO2 concentration. For live cell imaging experiments cells were imaged in Leibovitz L-15 (Thermofisher) medium supplemented with 10% FBS, penicillin (100 U/ml) and streptomycin (100 μg/ml).

G2-synchronization was achieved through a 24-h treatment with 100 nM Palbociclib (Selleckchem) followed by 12 h release into normal medium, as described in (Scott et al., 2020, Trotter & Hagan 2020).

Where indicated, cells were stained with 20 nM sirDNA (Spirochrome) following manufacture’s protocol, treated with 100 nM Taxol (Sigma-Aldrich) or nocodazole (55 nM, Sigma-Aldrich).

### Gene editing

For CCNB1 tagging, RPE-1 FRT/TO cells were transfected using 500 ng of a modified version of the PX466 “All-in-One” plasmid containing Cas9D10A-T2A-mRuby and gRNAs targeting CCNB1 (5’-ACCGTTTACTTTTAATAAAGCTTG-3’ and 5’-ACCGTAATATGTACAGATGGCACA-3’). The all-in-one plasmid was cotransfected with 500 ng of repair plasmid designed as a fusion of LINKER-mEmerald flanked by two 850 bp arms, homologous to the genomic region around the Cas9 cutting site. 72 hours post transfection, 50000 mRuby positive cells were sorted in a 1 cm well and expanded for one week before a second sorting of single cells in 96 well plates. The presence of mEmerald tag was identified through PCR using primers forward 5’-CAAATGCTTCTCCTATGTGACAGG-3’ and the reverse 5’-TTCAGGTGGGTGGGATTTAG-3’. PCR products of positive clones were sequenced using the same primers.

For Cdk1-mScarlet expression, RPE-1 FRT/TO CCNB1^+/+^ cell line was transfected with pcDNA5-FRT/TO-Cdk1-alpha-mScarlet and pOG44 (Invitrogen) using a 1:5 ratio. For mEm-alpha-mScarlet and mEmerald and mScarlet expression, RPE-1 FRT/TO cells were transfected with either pcDNA5-FRT/TO-mEm-alpha-mScarlet or pcDNA5-FRT/TO-mScarlet together with pOG44 using a 1:5 ratio. All transfections were followed by a two weeks selection using Geneticin (Gibco) 0.4 mg/ml. Gene expression was induced using tetracycline (Calbiochem) 1 μg/ml. In FCS experiments tetracycline was added 3 hours before imaging, in immunoprecipitation (Fig. S4C) tetracycline was added 16 hours before lysis.

All transfections were performed by electroporation using a Neon Transfection System (Invitrogen) with 2 pulses at 1400V for 20ms, using 1 μg total DNA per million cells.

### Protein extraction

In Fig S1, RPE cells were trypsinized and incubated in lysis buffer (150 mM NaCl, 50 mM Tris pH 7.4, 0.5% NP-40) supplemented with HALT protease/phosphatase inhibitor cocktail (Thermo Fisher Scientific) for 30 min at 4°C before clarification.

In Fig. 2, 4 and S4, following G2 Synchronization, RPE cells were trypsinized and resuspended in IP buffer (150 mM NaCl, 50 mM Tris pH 7.4, 2.5 mM MgCl2, 1 mM EGTA, 1 mM DTT, 1 mM PMSF) supplemented with HALT protease/phosphatase inhibitor cocktail. Cells were lysed through N2 cavitation, incubating the cells at 1500 psi, 20 min at 4°C and rapidly releasing the pressure

In all experiments, lysates were then clarified through centrifugation (14’000 g, 20 minutes, 4°C) and then quantified using Bradford Reagent (Bio-Rad Laboratories) according to manufacturer’s instructions.

### Immunodepletion & Immunoprecipitation

For Cdk1 immunodepletion (Fig. 2 and S3), 100 ug of clarified lysates were diluted to a final concentration of 2 ug/ul and incubated with 60 ul of Dynabeads prebound to 7.5 ug of either anti-Cdk1 Antibody (BD Biosciences - 610037) or Mouse Ig-G, in a total volume of 100 ul for 2 h at 4° C. The incubation with beads was repeated two (Fig 2A, S3A) or three times (Fig. 2D, S3D), depending on the experiment. 40 ug of either the immunodepleted lysates or the input were used for SDS-PAGE.

For Cyclin B1 immunoprecipitation (Fig. 2E, S3D, Fig 4D), 30 ul of Dynabeads were crosslinked to 3.75 ug of either anti-Cyclin B1 Antibody (GSN1 – SantaCruz – sc-245) or Mouse Ig-G, through incubation in 20 mM Dimethyl Pimelimidate solution, 20 min, RT. 200 ug of clarified lysates were then added to the beads and incubated for 3 h at 4° C.

For mEmerald and mScarlet immunoprecipitation (Fig. S4C), 30 ul of either GFP-Trap, RFP-Trap (Chromotek) or crosslinked IgG control (see above) magnetic beads were incubated 2 h at 4°C with 400 ug of clarified lysates diluted to a final concentration of 2 ug/ul.

For all IPs, beads were washed 5 times with IP buffer and then incubated 5 minutes at 65°C in 30 ul 2X Sample Loading Buffer, prior to SDS-PAGE.

### Immunoblotting

40 ug of RPE cell lysates were separated through SDS-PAGE on a 4–12% NuPAGE gel (Invitrogen) and transferred to an Immobilon-FL polyvinylidene fluoride membrane (IPFL00010, Millipore). The membrane was blocked with 5% Milk, 0.1 % Tween, PBS and incubated overnight with primary antibodies at 4°C in 2.5% Milk, 0.1 % Tween, PBS. The following day the membrane was washed with 0.1% Tween PBS and incubated with secondary antibodies in 2.5% Milk, 0.1 % Tween, PBS for 1h at RT.

Primary antibodies were used at the indicated concentrations: and anti-CCNB1 (1:1000, SantaCruz – GSN1, sc-245), anti-Cdk1 (1:1000, BD Biosciences - 610037), anti-Cdk2 (1:1000, 78B2 – Cell Signalling), anti-Tubulin (1:3000, ab6046 - Abcam). IRDye800CW donkey anti-mouse (926–32212, LI-COR), IRDye800CW donkey anti-rabbit (926–32213, LI-COR), IRDye680CW donkey anti-mouse (926–68072, LI-COR), and IRDye680CW donkey anti-rabbit (926–68073, LI-COR) secondary antibodies were all used at 1:10,000.

Proteins were visualized with LI-COR Odyssey CLx scanner (LI-COR Biosciences). Western blot quantification in Fig. 2 and S3 and 4D was performed calculating the area under the curve using Fiji’s “Gel” plugin. Values were adjusted by fitting to a straight-line function obtained by immunoblotting serial dilutions of protein lysate. In in Fig. 2 and S3 values were normalized to the input value. Regarding Fig. 4D, values were normalized on the 6 hour time point.

### Live-Cell Imaging

Mitotic time measurements (Fig. S1D) were obtained using differential interference contrast (DIC) imaging on a Nikon Eclipse microscope (Nikon) equipped with a 20× 0.75 NA objective (Nikon), a Flash 4.0 CMOS camera (Hamamatsu) and an analyser in the emission wheel for DIC imaging. Single plane images were taken every 3 minutes, for 24 hours using micromanager software (uManager) and analysed using FiJi (ImageJ).

Image series displayed in Fig. S1B and C were obtained on a Marianas confocal spinning-disk microscope system (Intelligent Imaging Innovations, Inc.) equipped with a laser stack for 445 nm/488 nm/514 nm/561 nm lasers, a 63× 1.2 NA objective (Carl Zeiss) and a Flash4 CMOS camera (Hamamatsu). 8 Z stacks (Step size = 1 μm) were taken every 30 s, for 90 minutes, using 20% 488 nm laser power and 10 % 647 nm laser power, for 50 ms exposure, using Slidebook 6 software (Intelligent Imaging Innovation, Inc.).

Widefield microscopy experiments (Fig. 4F, S5D) were performed using an Nikon Eclipse microscope (Nikon) equipped with a 40× 1.30 NA objective (Nikon) and a Flash 4.0 CMOS camera (Hamamatsu), recording 488 nm emission with 150 ms exposure. Single plane images were taken every 3 minutes, for 24 hours using micromanager software (uManager) and analysed using FiJi (ImageJ). Raw Integrated Density (RID) of the whole cell was normalized to the RID of the same cell 50 frames (150 min) before nuclear envelope breakdown. Multiple asynchronous cells were aligned on their NEBD time.

### Chromosome Spreads

For chromosome spreads (Fig. S2 E, D), following a 3-hours treatment with 100 ng/mL colcemid (GIBCO) cells were trypsinized and recovered in a falcon tube. Cell suspension was centrifuged for 3 minutes at 250 g and resuspended in 5 mL of 75 mM KCl, added dropwise. After a 15 minute-incubation at 37°C, 10 drops of Carnoys Fixative (3:1 methanol:acetic acid) were added. Following a 5 minute centrifugation at 200 g, the cell pellet was resuspended in 5 mL of Carnoys Fixative. After 90 min at −20°C, a second fixation was performed using 5 mL of Carnoys Fixative at room temperature for 15 minutes. Cells were then centrifuged at 200 g for 5 minutes, the supernatant removed, and the pellet resuspended in 200 μL of Carnoys Fixative.Spreads were performed by dropping the cells on wet slides in a wet chamber from 30-40 cm height. Chromosome spreads were aged at room temperature for 30 minutes, then incubated for 30 minutes with 1:50 Giemsa stain:Giemsa buffer, and finally washed in PBS pH 7.8. Once dried, slides were mounted with DPX mountant (Sigma-Aldrich) and coverslips and incubated overnight at room temperature.

Transmitted light images of metaphase spreads were captured using a 63× 1.4 NA lens on a Marianas confocal spinning-disk microscope system, and the number of chromosomes per cell was counted using ImageJ software.

### Flow Cytometry

In flow cytometry experiments (Fig. 4A, 5A) cells were detached, washed with PBS and fixed with 70% ethanol 4 hours −20°C. After fixation cells were stained 20 minutes in a 1 ug/ml Propidium Iodide (PI) solution, supplemented with 10 ug/ml RNAse (Sigma). Stained cells were acquired using a LSR II flow cytometers (BD bioscience). Cell cycle profiles were analysed using the software FlowJo.

### Calculating Cdk1-CycB binding

To calculate the fraction of Cyclin B1 engaged in Cdk1 binding over time, Cyclin B1 concentration increase was modelled by fitting FCS calculation (Fig. 4C) with a single exponential function (equation 3).

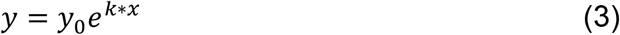

Where y_0_ = 5.561 and k = 0.04232, R = 0.8703. Cdk1 increase was modelled using a straight-line equation based on Fig. 4D and using initial Cdk1 concentration of 629 nM (Beck et al., 2011), slope =0.1509, y_0_ = 74.37, R = 0.9292.

### Data Analysis & Statistics

Statistical analysis, fitting and plotting were performed with Prism 8 (GraphPad). Graphs in fig. S2 were realized using following the “Superplots” pipeline and Python 3.7.0 (Lord et al., 2020).

### FCS instrumentation and measurements

The FCS and FCCS experiments were performed on a Leica TCS SP8 confocal microscope (DMI8; Leica). The samples were illuminated using a white light wavelength-adjustable pulsed laser that was focused to the back focal plane of a Leica HC PL APO CS2 63x/1.20 water immersion objective. The wavelengths used were 488 nm for samples expressing mEmerald or EGFP, and 569 nm for samples expressing mScarlet. The pinhole size was set to 1 airy unit and the emitted signal was recorded using Leica HyD SMD (single molecule detection) detectors with user-adjustable detection range. For mEmerald and EGFP, we used a detection range of 505-540 nm and for mScarlet a detection range of 580-625 nm.

Prior to each FCS/FCCS experiment, the objective’s collar was corrected to reduce aberrations, and the system was calibrated using Atto 488 and Atto 565 to determine the effective confocal volume and structure factor at 488 nm and 569 nm excitation respectively. The cells seeded on a μ-Slide 8 well ibiTreat dish were then measured for 10 s at 37 °C. The recorded signal was computed to generate auto- and cross-correlation functions and fit using Leica LAS X SMD FCS module. All measurements were fit with a three-dimensional free diffusion triplet (3D-triplet) model. The number of diffusing components in the fitting model were determined using Akaike information criterion (AIC) and F-tests in GraphPad Prism (Tsutsumi et al., 2016). We found that for CCNB1-mEm, a 3D two-component triplet model (3D-2particle-triplet model) is the most suitable model, whereas for freely diffusing EGFP in RPE cells, 3D one-component triplet model (3D-1particle-triplet model) is the more appropriate model.

In our FCCS experiments, we calculated the cross-correlation quotient q as the ratio of the CCF amplitude to the ACF amplitude. The q value is a measure of the amount of cross-correlation between the two species which is a representative of the fraction of molecules in complexes (Yavas et al., 2016). The dissociation constant (K_D_) for the interaction between Cyclin B1-mEmerald and Cdk1-mScarlet was calculated using equation 2. The concentrations of unbound Cyclin B1-mEmerald, unbound Cdk1-mScarlet and Cyclin B1-mEmerald-Cdk1-mScarlet complex for calculating K_D_ were estimated using FCCS as detailed in Veerapathiran et al., 2019 and Foo et al., 2012 (Foo et al., 2012; Veerapathiran et al., 2020).

## Author Contribution

M.B., L.C., S.V., C.C. and J.P. designed and interpreted the experiments and M.B., L.C., S.V., C.R., C. C. performed the experiments. M.B., L.C., S.V. and J.P. wrote the paper. J.P. conceived the study and acquired funding. Co-first authors appear in alphabetical order.

## Acknowledgements

We are grateful Andrew Harrison for pioneering the FCS work in our laboratory, and to all members of our lab for fruitful discussions. This work was supported by an Investigator Award from Wellcome (209470/Z/17/Z). We gratefully acknowledge the support of the ICR Core facilities, in particular light microscopy and flow cytometry.

## Supplementary Material

**Figure Supplementary 1:**
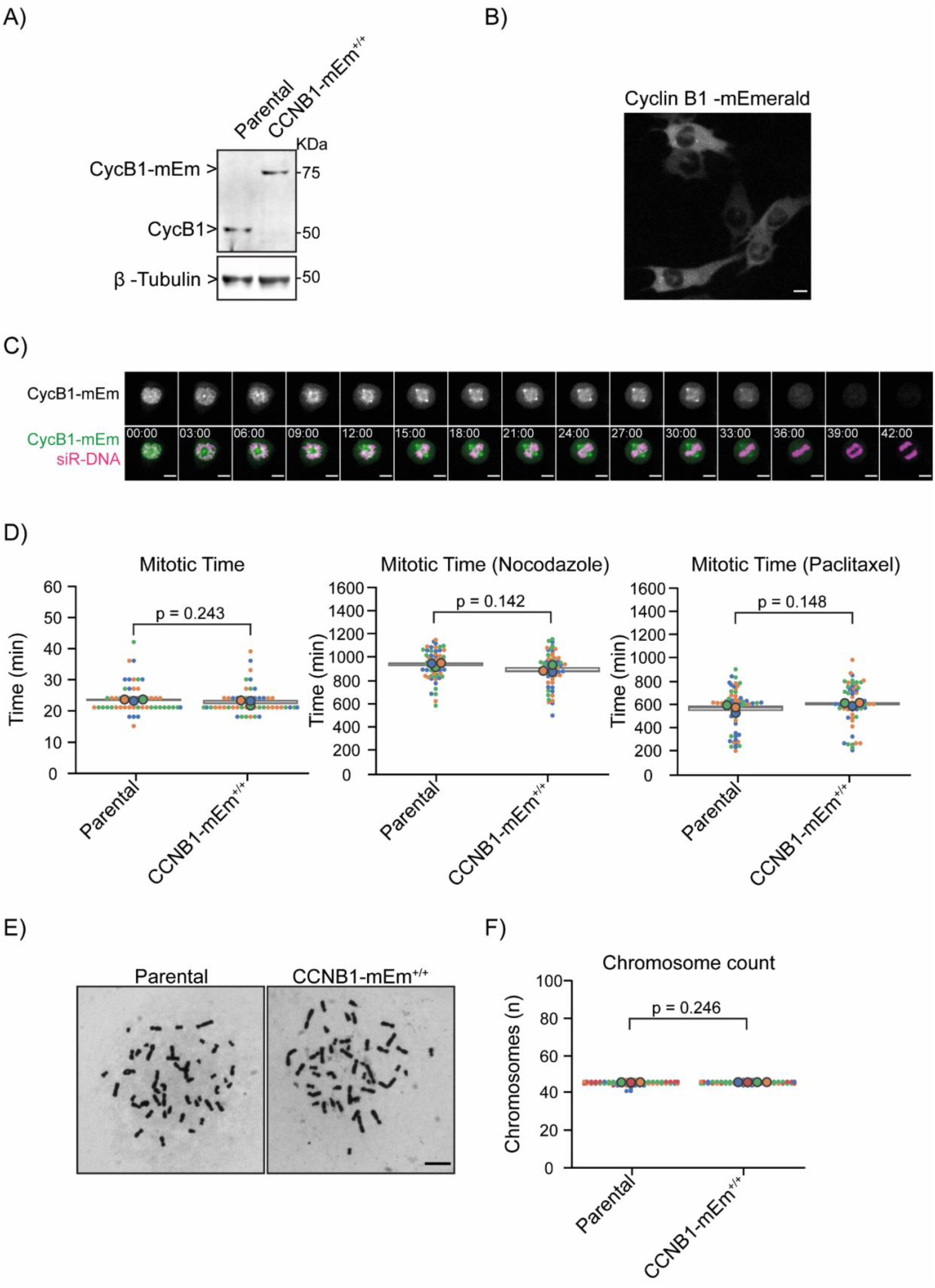
Characterization of RPE CCNB1-mEmerald^+/+^ cells. A) Anti-Cyclin B1 and anti-αTubulin immunoblots of lysates of parental RPE or RPE CCNB1-mEmerald^+/+^ cells showing that in the RPE CCNB1-mEmerald cells all Cyclin B1 molecules are tagged with mEmerald and are expressed at the same level as untagged Cyclin B1. Molecular mass indicated on the right. B) Representative fluorescence confocal image of RPE CCNB1-mEmerald^+/+^ cells in interphase. Scale bar corresponds to 20 μm. C) Representative fluorescence confocal images over time of a RPE CCNB1-mEmerald^+/+^ cells progressing through mitosis. Time is expressed as mm:ss. Scale bars correspond to 10 μm. D) Dot plots of the mitotic timing of parental RPE or RPE CCNB1-mEmerald^+/+^ cells, untreated (left), treated with 55 nM nocodazole (middle) or 100 nM paclitaxel (right). Each dot represents one cell. Statistical significance was determined using an unpaired t-test. E) Representative chromosome spreads of parental RPE and RPE CCNB1-mEmerald^+/+^ cells. Scale bar corresponds to 20 μm. F) Dot plots of the chromosome number of parental RPE or RPE CCNB1-mEmerald^+/+^ cells. Each dot represents one cell. Statistical significance was determined using unpaired t-test with Welch’s correction. In all plots the large dots represent the median of independent experiments. N = 3 independent experiments.

**Figure Supplementary 2:**
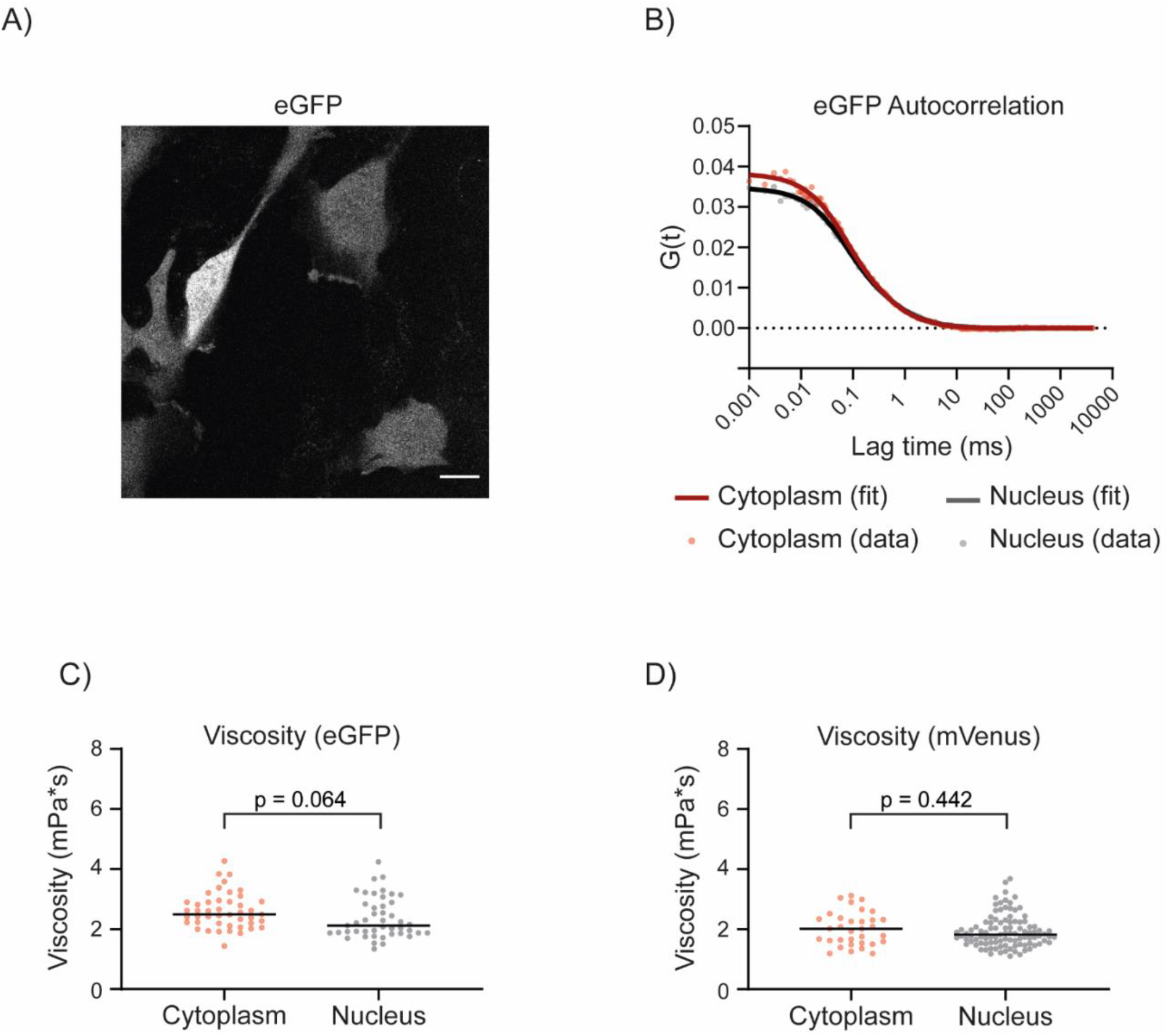
Viscosity of RPE cells. A) Representative fluorescence confocal image of RPE cells expressing eGFP. Scale bar corresponds to 20 μm. B) Graph representing the autocorrelation function of eGFP over time. C) Dot plots representing the viscosity in the nucleus and the cytoplasm of RPE cells expressing eGFP. Horizontal bars indicate medians, each dot corresponds to a single FCS measurement. D) Dot plots representing the viscosity in the nucleus and the cytoplasm of RPE cells expressing mVenus. Horizontal bars indicate medians, each dot corresponds to a single FCS measurement. Statistical significance was determined using an unpaired t-test. N = 3 independent experiments.

**Figure Supplementary 3:**
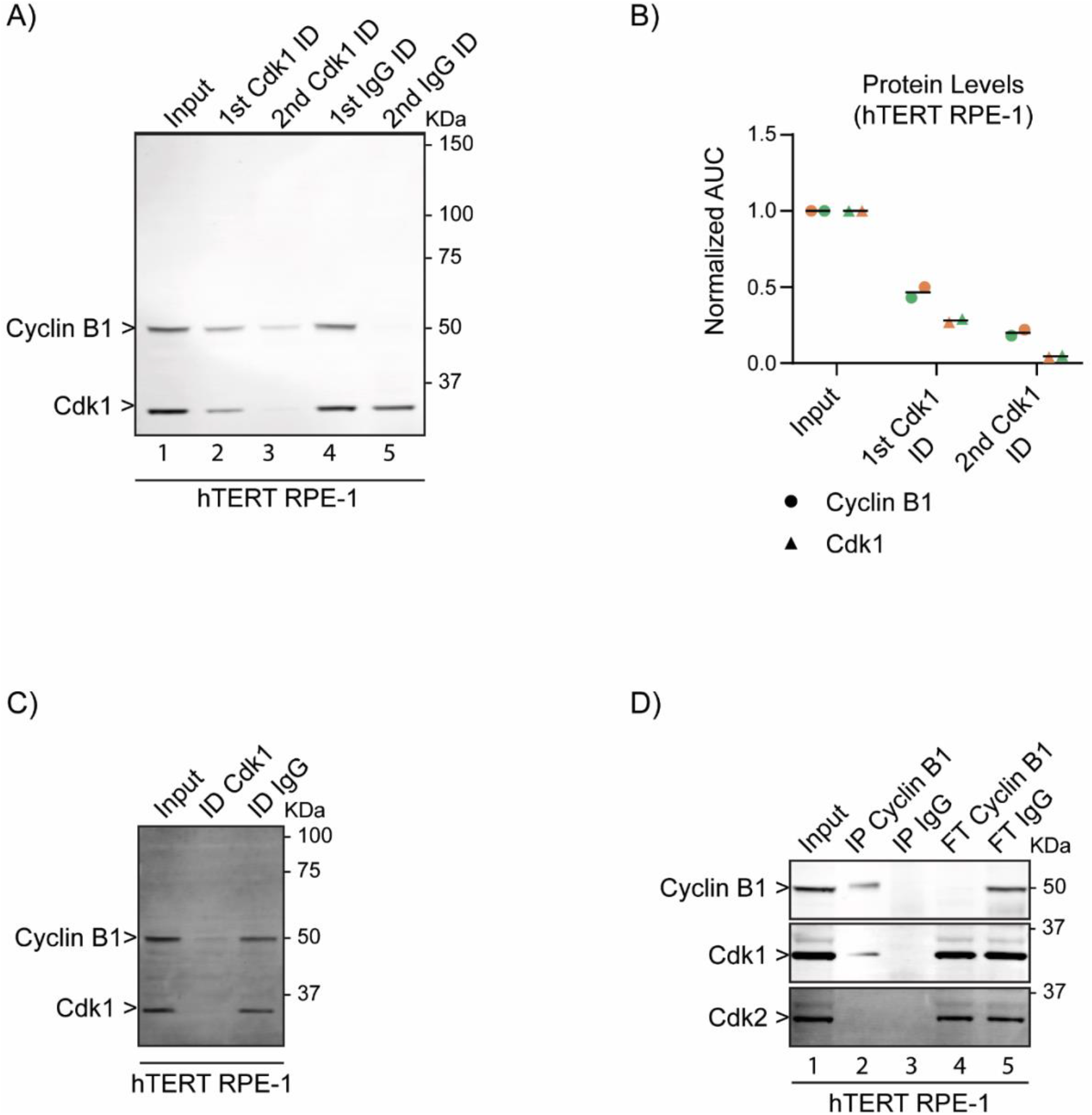
A fraction of Cyclin B1 is not in complex with Cdk1 in RPE cells during G2 phase. A) Anti-Cyclin B1 and anti-Cdk1 immunoblot of G2 phase cell lysates before (1st) and after (2nd and 3rd lanes) immunodepleting Cdk1, compared with control immunodepletion with IgG (4th and 5th lanes). (Note that we do not have a definitive explanation for the apparent depletion of Cyclin B1 on control beads in the 2nd depletion.) B) Quantification of Cyclin B1 and Cdk1 levels measured by immunoblotting, before and after immunodepletion of Cdk1. C) Anti-Cyclin B1 and anti-Cdk1 immunoblot of G2 phase cell lysates before and after immunodepletion of Cdk1 to at or below detectable levels, or after control immunodepletion. D) Anti-Cyclin B1, anti-Cdk1 and anti-Cdk2 immunoblots of G2 phase cell lysates before (1^st^ lane) and after immunoprecipitation with anti-Cyclin B1 antibody (2^nd^ lane), or immunoprecipitation with control anti-IgG antibody (3^rd^ lane) and the unbound fractions from the respective immunodepletions (4^th^ and 5^th^ lanes). For all graphs, individual dots represent biological replicates, horizontal lines indicate medians. For all panels, N=2 independent experiments. ID= immunodepletion, IP = immunoprecipitation, FT = Flow Through. Molecular mass indicated for all gels on the right.

**Figure Supplementary 4:**
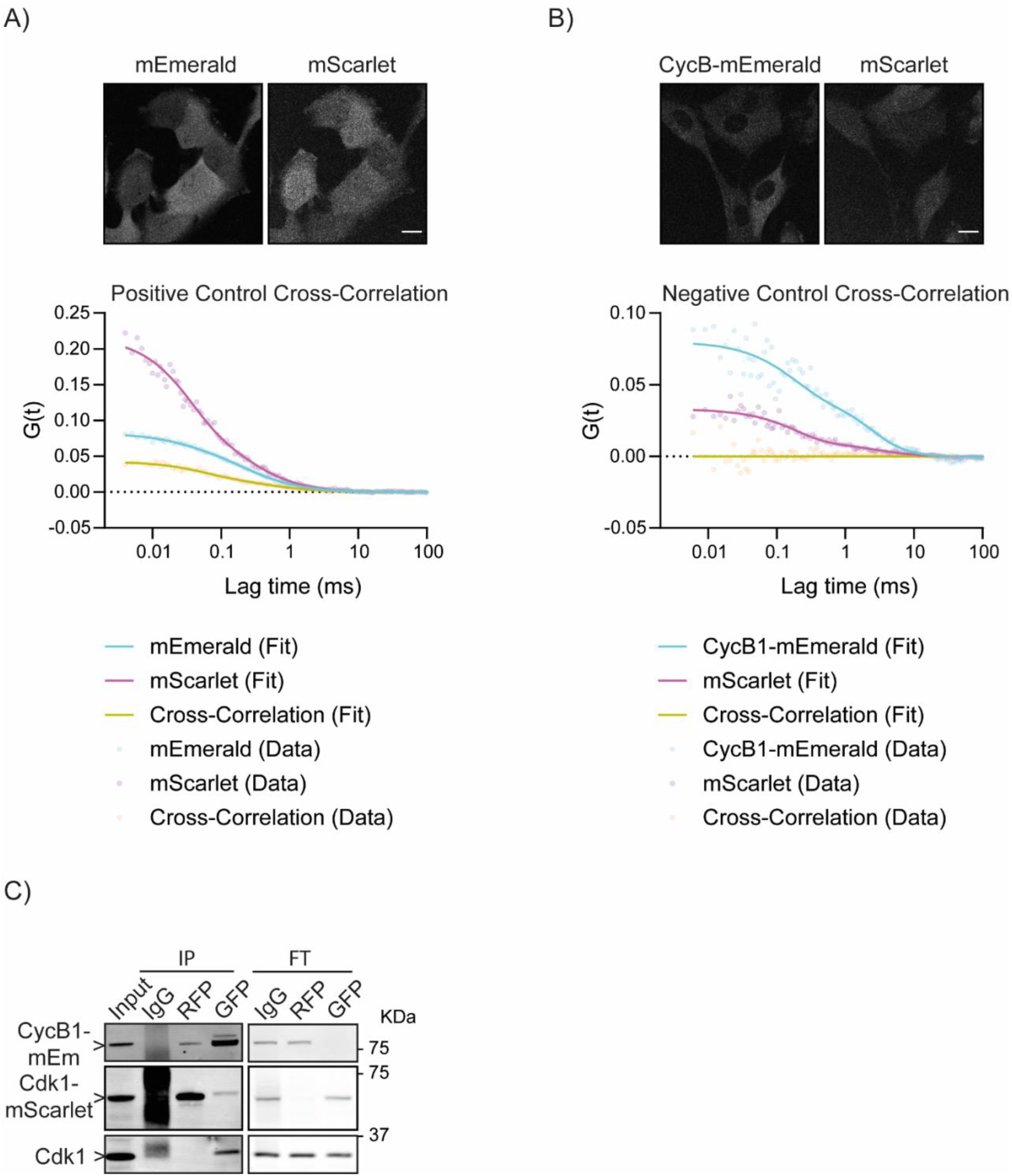
FCCS controls. A) Top panel: representative fluorescence confocal images of RPE cells expressing an mEmerald-mScarlet fusion protein. Scale bar corresponds to 20 μm. Bottom panel: graph of the autocorrelation function of mEmerald and mScarlet and the cross-correlation function between the two. Note the positive cross-correlation signal, as expected. B) Top panel: representative fluorescence confocal images of RPE CCNB1-mEmerald^+/+^ cells expressing mScarlet. Scale bar corresponds to 20 μm. Bottom panel: graph of the autocorrelation function of Cyclin B1-mEmerald and mScarlet and the cross-correlation function between the two. Note the lack of cross correlation between the molecules. C) Anti-Cyclin B1 and anti-Cdk1 immunoblots of RPE CCNB1-mEmerald^+/+^ cells lysates before (1^st^ lane) and after immunoprecipitation with anti-IgG antibody (2^nd^ lane), anti-RFP antibody (3^rd^ lane), anti-GFP antibody (4^th^ lane) and lysates following the respective immunodepletions (5^th^, 6^th^ and 7^th^ lanes). Note that Cyclin B1-mEmerald co-precipitates with Cdk1-mScarlet. Molecular mass indicated on the right. N=1 experiment. For all FCCS experiments N=2 or more.

**Figure Supplementary 5:**
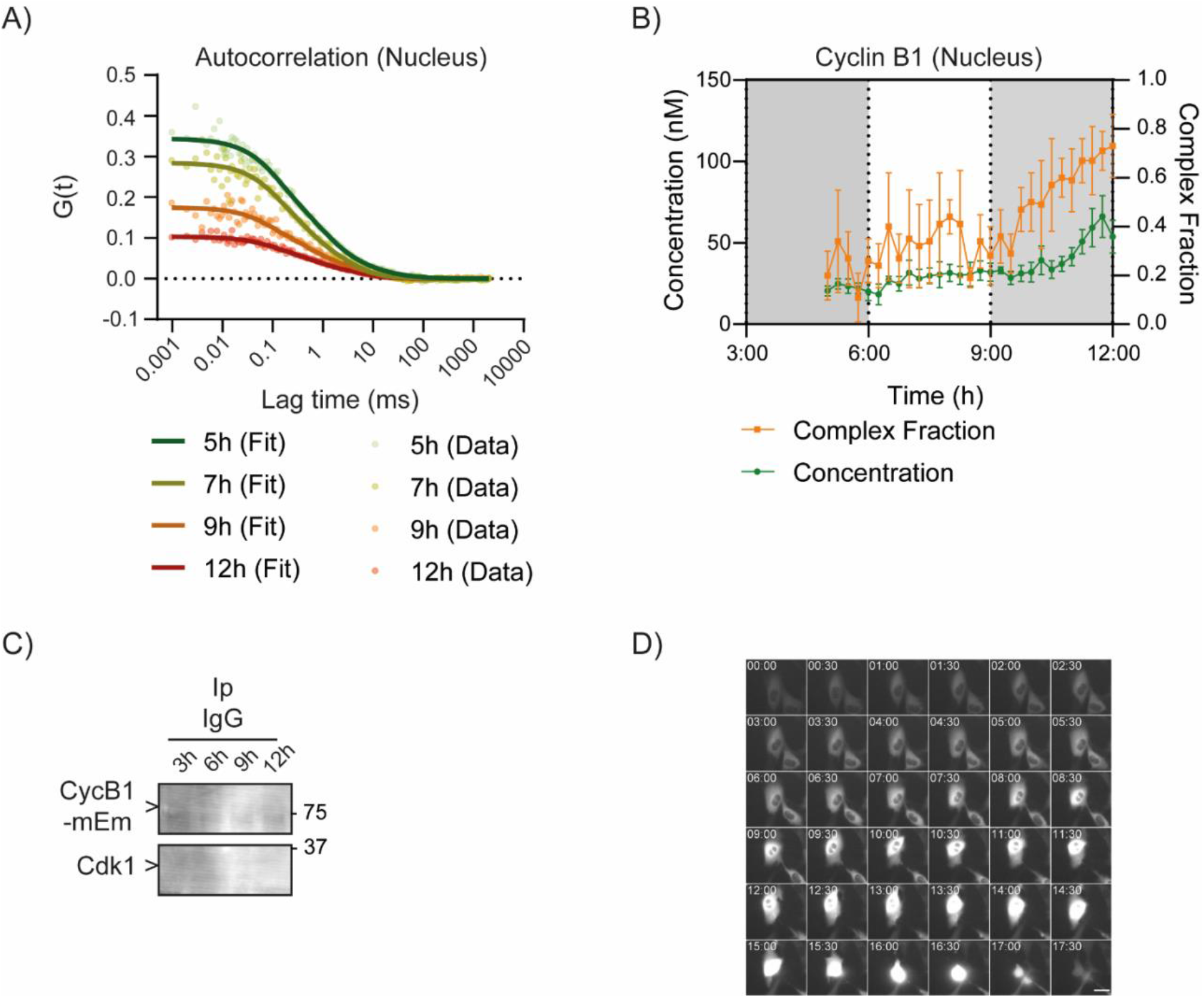
Time resolved Cyclin B1 FCS. A) Graph representing the FCS autocorrelation functions of Cyclin B1-mEmerald at different the indicated time points after release from Palbociclib-arrest. Each dot represents the measurement from one cell. B) Graph showing time resolved Cyclin B1-mEmerald concentration (left axis) and the fraction of Cyclin B1 in complex with Cdk1 (right axis) estimated from FCS measurements in the nucleus. C) anti-Cyclin B1 and anti-Cdk1 immuno-blot of synchronized RPE CCNB1-mEmerald^+/+^ cells lysates after immunoprecipitation with anti-IgG antibody. D) Representative widefield images over time of a RPE CCNB1-mEmerald^+/+^ cells. Time is expressed at hh:mm. Scale bars correspond to 10 μm. For all panels, N=2 independent experiments, time indicates hours after Palbociclib release.

## References

Arooz, T., Yam, C.H., Siu, W.Y., Lau, A., Li, K.K., Poon, R.Y., 2000. On the concentrations of cyclins and cyclind-ependent kinases in extracts of cultured human cells. Biochemistry 39, 9494–9501. https://doi.org/10.1021/bi0009643

Bacia, K., Kim, S.A., Schwille, P., 2006. Fluorescence cross-correlation spectroscopy in living cells. Nat Methods 3, 83–89. https://doi.org/10.1038/nmeth822

Balleza, E., Kim, J.M., Cluzel, P., 2018. Systematic characterization of maturation time of fluorescent proteins in living cells. Nat Methods 15, 47–51. https://doi.org/10.1038/nmeth.4509

Becher, I., Andrés-Pons, A., Romanov, N., Stein, F., Schramm, M., Baudin, F., Helm, D., Kurzawa, N., Mateus, A., Mackmull, M.-T., Typas, A., Müller, C.W., Bork, P., Beck, M., Savitski, M.M., 2018. Pervasive Protein Thermal Stability Variation during the Cell Cycle. Cell 173, 1495–1507.e18. https://doi.org/10.1016/j.cell.2018.03.053

Beck, M., Schmidt, A., Malmstroem, J., Claassen, M., Ori, A., Szymborska, A., Herzog, F., Rinner, O., Ellenberg, J., Aebersold, R., 2011. The quantitative proteome of a human cell line. Mol Syst Biol 7, 549. https://doi.org/10.1038/msb.2011.82

Bindels, D.S., Haarbosch, L., van Weeren, L., Postma, M., Wiese, K.E., Mastop, M., Aumonier, S., Gotthard, G., Royant, A., Hink, M.A., Gadella, T.W.J., 2017. mScarlet: a bright monomeric red fluorescent protein for cellular imaging. Nat. Methods 14, 53–56. https://doi.org/10.1038/nmeth.4074

Brock, R., Jovin, T.M., 1998. Fluorescence correlation microscopy (FCM)-fluorescence correlation spectroscopy (FCS) taken into the cell. Cell Mol Biol (Noisy-le-grand) 44, 847–856.

Brown, N.R., Korolchuk, S., Martin, M.P., Stanley, W.A., Moukhametzianov, R., Noble, M.E.M., Endicott, J.A., 2015. CDK1 structures reveal conserved and unique features of the essential cell cycle CDK. Nat Commun 6, 6769. https://doi.org/10.1038/ncomms7769

Cheng, A., Kaldis, P., Solomon, M.J., 2000. Dephosphorylation of Human Cyclin-dependent Kinases by Protein Phosphatase Type 2Cα and β2 Isoforms *. Journal of Biological Chemistry 275, 34744–34749. https://doi.org/10.1074/jbc.M006210200

Clute, P., Pines, J., 1999. Temporal and spatial control of cyclin B1 destruction in metaphase. Nat Cell Biol 1, 82–87. https://doi.org/10.1038/10049

Coulonval, K., Kooken, H., Roger, P.P., 2011. Coupling of T161 and T14 phosphorylations protects cyclin B-CDK1 from premature activation. Mol Biol Cell 22, 3971–3985. https://doi.org/10.1091/mbc.E11-02-0136

Cubitt, A.B., Woollenweber, L.A., Heim, R., 1998. Chapter 2: Understanding Structure—Function Relationships in the Aequorea victoria Green Fluorescent Protein, in: Sullivan, K.F., Kay, S.A. (Eds.), Methods in Cell Biology, Green Fluorescent Proteins. Academic Press, pp. 19–30. https://doi.org/10.1016/S0091-679X(08)61946-9

Darbon, J.M., Devault, A., Taviaux, S., Fesquet, D., Martinez, A.M., Galas, S., Cavadore, J.C., Dorée, M., Blanchard, J.M., 1994. Cloning, expression and subcellular localization of the human homolog of p40MO15 catalytic subunit of cdk-activating kinase. Oncogene 9, 3127–3138.

Deota, S., Rathnachalam, S., Namrata, K., Boob, M., Fulzele, A., Radhika, S., Ganguli, S., Balaji, C., Kaypee, S., Vishwakarma, K.K., Kundu, T.K., Bhandari, R., Gonzalez de Peredo, A., Mishra, M., Venkatramani, R., Kolthur-Seetharam, U., 2019. Allosteric Regulation of Cyclin-B Binding by the Charge State of Catalytic Lysine in CDK1 Is Essential for Cell-Cycle Progression. J Mol Biol 431, 2127–2142. https://doi.org/10.1016/j.jmb.2019.04.005

Desai, D., Wessling, H.C., Fisher, R.P., Morgan, D.O., 1995. Effects of phosphorylation by CAK on cyclin binding by CDC2 and CDK2. Molecular and Cellular Biology 15, 345. https://doi.org/10.1128/mcb.15.1.345

Doudna, J.A., Charpentier, E., 2014. Genome editing. The new frontier of genome engineering with CRISPR-Cas9. Science 346, 1258096. https://doi.org/10.1126/science.1258096

Draetta, G., Brizuela, L., Potashkin, J., Beach, D., 1987. Identification of p34 and p13, human homologs of the cell cycle regulators of fission yeast encoded by cdc2+ and suc1+. Cell 50, 319–325. https://doi.org/10.1016/0092-8674(87)90227-3

Ducommun, B., Brambilla, P., Félix, M.A., Franza, B.R., Karsenti, E., Draetta, G., 1991. cdc2 phosphorylation is required for its interaction with cyclin. EMBO J 10, 3311–3319.

Dunsing, V., Luckner, M., Zühlke, B., Petazzi, R.A., Herrmann, A., Chiantia, S., 2018. Optimal fluorescent protein tags for quantifying protein oligomerization in living cells. Sci Rep 8, 10634. https://doi.org/10.1038/s41598-018-28858-0

Enderlein, J., Gregor, I., Patra, D., Dertinger, T., Kaupp, U.B., 2005. Performance of Fluorescence Correlation Spectroscopy for Measuring Diffusion and Concentration. ChemPhysChem 6, 2324–2336. https://doi.org/10.1002/cphc.200500414

Fesquet, D., Labbé, J.C., Derancourt, J., Capony, J.P., Galas, S., Girard, F., Lorca, T., Shuttleworth, J., Dorée, M., Cavadore, J.C., 1993. The MO15 gene encodes the catalytic subunit of a protein kinase that activates cdc2 and other cyclin-dependent kinases (CDKs) through phosphorylation of Thr161 and its homologues. The EMBO Journal 12, 3111.

Fisher, R.P., Morgan, D.O., 1994. A novel cyclin associates with MO15/CDK7 to form the CDK-activating kinase. Cell 78, 713–724. https://doi.org/10.1016/0092-8674(94)90535-5

Fleming, P.J., Fleming, K.G., 2018. HullRad: Fast Calculations of Folded and Disordered Protein and Nucleic Acid Hydrodynamic Properties. Biophysical Journal 114, 856–869. https://doi.org/10.1016/j.bpj.2018.01.002

Foo, Y.H., Naredi-Rainer, N., Lamb, D.C., Ahmed, S., Wohland, T., 2012. Factors affecting the quantification of biomolecular interactions by fluorescence cross-correlation spectroscopy. Biophys J 102, 1174–1183. https://doi.org/10.1016/j.bpj.2012.01.040

Frisa, P.S., Jacobberger, J.W., 2009. Cell Cycle-Related Cyclin B1 Quantification. PLoS ONE 4, e7064. https://doi.org/10.1371/journal.pone.0007064

Gould, K.L., Moreno, S., Owen, D.J., Sazer, S., Nurse, P., 1991. Phosphorylation at Thr167 is required for Schizosaccharomyces pombe p34cdc2 function. EMBO J 10, 3297–3309.

Gyuris, J., Golemis, E., Chertkov, H., Brent, R., 1993. Cdi1, a human G1 and S phase protein phosphatase that associates with Cdk2. Cell. https://doi.org/10.1016/0092-8674(93)90498-F

Hagting, A., Karlsson, C., Clute, P., Jackman, M., Pines, J., 1998. MPF localization is controlled by nuclear export. EMBO J 17, 4127–4138. https://doi.org/10.1093/emboj/17.14.4127

Hannon, G.J., Casso, D., Beach, D., 1994. KAP: a dual specificity phosphatase that interacts with cyclin-dependent kinases. PNAS 91, 1731–1735. https://doi.org/10.1073/pnas.91.5.1731

Hillesheim, L.N., Chen, Y., Müller, J.D., 2006. Dual-Color Photon Counting Histogram Analysis of mRFP1 and EGFP in Living Cells. Biophys J 91, 4273–4284. https://doi.org/10.1529/biophysj.106.085845

Hink, M.A., Griep, R.A., Borst, J.W., van Hoek, A., Eppink, M.H., Schots, A., Visser, A.J., 2000. Structural dynamics of green fluorescent protein alone and fused with a single chain Fv protein. J Biol Chem 275, 17556–17560. https://doi.org/10.1074/jbc.M001348200

Jackman, M., Lindon, C., Nigg, E.A., Pines, J., 2003. Active cyclin B1-Cdk1 first appears on centrosomes in prophase. Nat Cell Biol 5, 143–148. https://doi.org/10.1038/ncb918

Jackman, M., Marcozzi, C., Barbiero, M., Pardo, M., Yu, L., Tyson, A.L., Choudhary, J.S., Pines, J., 2020. Cyclin B1-Cdk1 facilitates MAD1 release from the nuclear pore to ensure a robust spindle checkpoint. J Cell Biol 219, e201907082. https://doi.org/10.1083/jcb.201907082

Kaldis, P., Ojala, P.M., Tong, L., Mäkelä, T.P., Solomon, M.J., 2001. CAK-independent activation of CDK6 by a viral cyclin. Mol Biol Cell 12, 3987–3999. https://doi.org/10.1091/mbc.12.12.3987

Karuna, A., Masia, F., Chappell, S., Errington, R., Hartley, A.M., Jones, D.D., Borri, P., Langbein, W., 2020. Quantitative Imaging of B1 Cyclin Expression Across the Cell Cycle Using Green Fluorescent Protein Tagging and Epifluorescence. Cytometry A 97, 1066–1072. https://doi.org/10.1002/cyto.a.24038

Kim, S.A., Heinze, K.G., Schwille, P., 2007. Fluorescence correlation spectroscopy in living cells. Nat Methods 4, 963–973. https://doi.org/10.1038/nmeth1104

Kõivomägi, M., Ord, M., Iofik, A., Valk, E., Venta, R., Faustova, I., Kivi, R., Balog, E.R.M., Rubin, S.M., Loog, M., 2013. Multisite phosphorylation networks as signal processors for Cdk1. Nat Struct Mol Biol 20, 1415–1424. https://doi.org/10.1038/nsmb.2706

Kõivomägi, M., Valk, E., Venta, R., Iofik, A., Lepiku, M., Morgan, D.O., Loog, M., 2011. Dynamics of Cdk1 substrate specificity during the cell cycle. Mol Cell 42, 610–623. https://doi.org/10.1016/j.molcel.2011.05.016

Krichevsky, O., Bonnet, G., 2002. Fluorescence correlation spectroscopy: the technique and its applications. Rep. Prog. Phys. 65, 251–297. https://doi.org/10.1088/0034-4885/65/2/203

Larochelle, S., Merrick, K.A., Terret, M.-E., Wohlbold, L., Barboza, N.M., Zhang, C., Shokat, K.M., Jallepalli, P.V., Fisher, R.P., 2007. Requirements for Cdk7 in the assembly of Cdk1/cyclin B and activation of Cdk2 revealed by chemical genetics in human cells. Mol Cell 25, 839–850. https://doi.org/10.1016/j.molcel.2007.02.003

Lee, K.M., Saiz, J.E., Barton, W.A., Fisher, R.P., 1999. Cdc2 activation in fission yeast depends on Mcs6 and Csk1, two partially redundant Cdk-activating kinases (CAKs). Curr Biol 9, 441–444. https://doi.org/10.1016/s0960-9822(99)80194-8

Lipinski, C., Hopkins, A., 2004. Navigating chemical space for biology and medicine. Nature 432, 855–861. https://doi.org/10.1038/nature03193

Liu, Y., Wu, C., Galaktionov, K., 2004. p42, a novel cyclin-dependent kinase-activating kinase in mammalian cells. J Biol Chem 279, 4507–4514. https://doi.org/10.1074/jbc.M309995200

Lord, S.J., Velle, K.B., Mullins, R.D., Fritz-Laylin, L.K., 2020. SuperPlots: Communicating reproducibility and variability in cell biology. J Cell Biol 219, e202001064. https://doi.org/10.1083/jcb.202001064

Luo, Q., Sewalt, E., Borst, J.W., Westphal, A.H., Boom, R.M., Janssen, A.E.M., 2019. Analysis and modeling of enhanced green fluorescent protein diffusivity in whey protein gels. Food Res Int 120, 449–455. https://doi.org/10.1016/j.foodres.2018.10.087

Ly, T., Whigham, A., Clarke, R., Brenes-Murillo, A.J., Estes, B., Madhessian, D., Lundberg, E., Wadsworth, P., Lamond, A.I., 2017. Proteomic analysis of cell cycle progression in asynchronous cultures, including mitotic subphases, using PRIMMUS. eLife 6, e27574. https://doi.org/10.7554/eLife.27574

Magde, D., Elson, E.L., Webb, W.W., 1974. Fluorescence correlation spectroscopy. II. An experimental realization. Biopolymers 13, 29–61. https://doi.org/10.1002/bip.1974.360130103

Morgan, D.O., 1997. Cyclin-dependent kinases: engines, clocks, and microprocessors. Annu Rev Cell Dev Biol 13, 261–291. https://doi.org/10.1146/annurev.cellbio.13.1.261

Murray, A.W., Kirschner, M.W., 1989. Cyclin synthesis drives the early embryonic cell cycle. Nature 339, 275–280. https://doi.org/10.1038/339275a0

Murray, A.W., Solomon, M.J., Kirschner, M.W., 1989. The role of cyclin synthesis and degradation in the control of maturation promoting factor activity. Nature 339, 280–286. https://doi.org/10.1038/339280a0

Nalepa, G., Barnholtz-Sloan, J., Enzor, R., Dey, D., He, Y., Gehlhausen, J.R., Lehmann, A.S., Park, S.-J., Yang, Y., Yang, X., Chen, S., Guan, X., Chen, Y., Renbarger, J., Yang, F.-C., Parada, L.F., Clapp, W., 2013. The tumor suppressor CDKN3 controls mitosis. J Cell Biol 201, 997–1012. https://doi.org/10.1083/jcb.201205125

Norbury, C., Blow, J., Nurse, P., 1991. Regulatory phosphorylation of the p34cdc2 protein kinase in vertebrates. EMBO J 10, 3321–3329.

Nurse, P., 1990. Universal control mechanism regulating onset of M-phase. Nature 344, 503–508. https://doi.org/10.1038/344503a0

Olsen, J.V., Vermeulen, M., Santamaria, A., Kumar, C., Miller, M.L., Jensen, L.J., Gnad, F., Cox, J., Jensen, T.S., Nigg, E.A., Brunak, S., Mann, M., 2010. Quantitative Phosphoproteomics Reveals Widespread Full Phosphorylation Site Occupancy During Mitosis. Science Signaling. https://doi.org/10.1126/scisignal.2000475

Patra, D., Wang, S.X., Kumagai, A., Dunphy, W.G., 1999. The Xenopus Suc1/Cks Protein Promotes the Phosphorylation of G2/M Regulators*. Journal of Biological Chemistry 274, 36839–36842. https://doi.org/10.1074/jbc.274.52.36839

Pines, J., 1996. Cell cycle: reaching for a role for the Cks proteins. Curr Biol 6, 1399–1402. https://doi.org/10.1016/s0960-9822(96)00741-5

Pines, J., Hagan, I., 2011. The Renaissance or the cuckoo clock. Philosophical Transactions of the Royal Society B: Biological Sciences 366, 3625–3634. https://doi.org/10.1098/rstb.2011.0080

Pines, J., Hunter, T., 1991. Human cyclins A and B1 are differentially located in the cell and undergo cell cycle-dependent nuclear transport. J Cell Biol 115, 1–17. https://doi.org/10.1083/jcb.115.1.1

Pines, J., Hunter, T., 1989. Isolation of a human cyclin cDNA: evidence for cyclin mRNA and protein regulation in the cell cycle and for interaction with p34cdc2. Cell 58, 833–846. https://doi.org/10.1016/0092-8674(89)90936-7

Poon, R.Y., Hunter, T., 1995. Dephosphorylation of Cdk2 Thr160 by the cyclin-dependent kinase-interacting phosphatase KAP in the absence of cyclin. Science 270, 90–93. https://doi.org/10.1126/science.270.5233.90

Poon, R.Y., Yamashita, K., Adamczewski, J.P., Hunt, T., Shuttleworth, J., 1993. The cdc2-related protein p40MO15 is the catalytic subunit of a protein kinase that can activate p33cdk2 and p34cdc2. EMBO J 12, 3123–3132.

Richardson, H.E., Stueland, C.S., Thomas, J., Russell, P., Reed, S.I., 1990. Human cDNAs encoding homologs of the small p34Cdc28/Cdc2-associated protein of Saccharomyces cerevisiae and Schizosaccharomyces pombe. Genes Dev. 4, 1332–1344. https://doi.org/10.1101/gad.4.8.1332

Ries, J., Yu, S.R., Burkhardt, M., Brand, M., Schwille, P., 2009. Modular scanning FCS quantifies receptor-ligand interactions in living multicellular organisms. Nat Methods 6, 643–645. https://doi.org/10.1038/nmeth.1355

Schenk, A., Ivanchenko, S., Röcker, C., Wiedenmann, J., Nienhaus, G.U., 2004. Photodynamics of Red Fluorescent Proteins Studied by Fluorescence Correlation Spectroscopy. Biophysical Journal 86, 384. https://doi.org/10.1016/S0006-3495(04)74114-4

Schwille, P., Meyer-Almes, F.J., Rigler, R., 1997. Dual-color fluorescence cross-correlation spectroscopy for multicomponent diffusional analysis in solution. Biophys J 72, 1878–1886. https://doi.org/10.1016/S0006-3495(97)78833-7

Scott, S.J., Suvarna, K.S., D’Avino, P.P., 2020. Synchronization of human retinal pigment epithelial-1 cells in mitosis. J Cell Sci 133. https://doi.org/10.1242/jcs.247940

Shen, B., Zhang, W., Zhang, J., Zhou, J., Wang, J., Chen, L., Wang, L., Hodgkins, A., Iyer, V., Huang, X., Skarnes, W.C., 2014. Efficient genome modification by CRISPR-Cas9 nickase with minimal off-target effects. Nat Methods 11, 399–402. https://doi.org/10.1038/nmeth.2857

Smedt, V.D., Poulhe, R., Cayla, X., Dessauge, F., Karaiskou, A., Jessus, C., Ozon, R., 2002. Thr-161 Phosphorylation of Monomeric Cdc2: REGULATION BY PROTEIN PHOSPHATASE 2C IN XENOPUSOOCYTES *. Journal of Biological Chemistry 277, 28592–28600. https://doi.org/10.1074/jbc.M202742200

Solomon, M.J., Harper, J.W., Shuttleworth, J., 1993. CAK, the p34cdc2 activating kinase, contains a protein identical or closely related to p40MO15. EMBO J 12, 3133–3142.

Strauss, B., Harrison, A., Coelho, P.A., Yata, K., Zernicka-Goetz, M., Pines, J., 2017. Cyclin B1 is essential for mitosis in mouse embryos, and its nuclear export sets the time for mitosis. Journal of Cell Biology 217, 179–193. https://doi.org/10.1083/jcb.201612147

Tassan, J.P., Schultz, S.J., Bartek, J., Nigg, E.A., 1994. Cell cycle analysis of the activity, subcellular localization, and subunit composition of human CAK (CDK-activating kinase). J Cell Biol 127, 467–478. https://doi.org/10.1083/jcb.127.2.467

Thomas, N., Kenrick, M., Giesler, T., Kiser, G., Tinkler, H., Stubbs, S., 2005. Characterization and gene expression profiling of a stable cell line expressing a cell cycle GFP sensor. Cell Cycle 4, 191–195. https://doi.org/10.4161/cc.4.1.1405

Trotter, E.W., Hagan, I.M., 2020. Release from cell cycle arrest with Cdk4/6 inhibitors generates highly synchronized cell cycle progression in human cell culture. Open Biol. 10, 200200. https://doi.org/10.1098/rsob.200200

Tsutsumi, M., Muto, H., Myoba, S., Kimoto, M., Kitamura, A., Kamiya, M., Kikukawa, T., Takiya, S., Demura, M., Kawano, K., Kinjo, M., Aizawa, T., 2016. In vivo fluorescence correlation spectroscopy analyses of FMBP-1, a silkworm transcription factor. FEBS Open Bio 6, 106–125. https://doi.org/10.1002/2211-5463.12026

Veerapathiran, S., Teh, C., Zhu, S., Kartigayen, I., Korzh, V., Matsudaira, P.T., Wohland, T., 2020. Wnt3 distribution in the zebrafish brain is determined by expression, diffusion and multiple molecular interactions. eLife 9, e59489. https://doi.org/10.7554/eLife.59489

Wachsmuth, M., Conrad, C., Bulkescher, J., Koch, B., Mahen, R., Isokane, M., Pepperkok, R., Ellenberg, J., 2015. High-throughput fluorescence correlation spectroscopy enables analysis of proteome dynamics in living cells. Nat Biotechnol 33, 384–389. https://doi.org/10.1038/nbt.3146

Walther, N., Hossain, M.J., Politi, A.Z., Koch, B., Kueblbeck, M., Ødegård-Fougner, Ø., Lampe, M., Ellenberg, J., 2018. A quantitative map of human Condensins provides new insights into mitotic chromosome architecture. Journal of Cell Biology 217, 2309–2328. https://doi.org/10.1083/jcb.201801048

Welch, P.J., Wang, J.Y., 1992. Coordinated synthesis and degradation of cdc2 in the mammalian cell cycle. PNAS 89, 3093–3097. https://doi.org/10.1073/pnas.89.7.3093

Xu, N., Chang, D.C., 2007. Different thresholds of MPF inactivation are responsible for controlling different mitotic events in mammalian cell division. Cell Cycle 6, 1639–1645. https://doi.org/10.4161/cc.6.13.4385

Yu, L., Lei, Y., Ma, Y., Liu, M., Zheng, J., Dan, D., Gao, P., 2021. A Comprehensive Review of Fluorescence Correlation Spectroscopy. Front. Phys. 0. https://doi.org/10.3389/fphy.2021.644450

